# Patho-transcriptomic analysis of invasive mucinous adenocarcinoma of the lung (IMA): comparison with lung adenocarcinoma with signet ring cell features (SRCC)

**DOI:** 10.1101/2024.06.13.598839

**Authors:** William D. Stuart, Masaoki Ito, Iris-Fink Baldauf, Takuya Fukazawa, Tomoki Yamatsuji, Tomoshi Tsuchiya, Hideo Watanabe, Morihito Okada, Eric L. Snyder, Mari Mino-Kenudson, Minzhe Guo, Yutaka Maeda

## Abstract

**Background:** Invasive mucinous adenocarcinoma (IMA) comprises ∼5% of lung adenocarcinoma. There is no effective therapy for IMA when surgical resection is not possible. IMA is sometimes confused with adenocarcinoma with signet ring cell features (SRCC) pathologically since both adenocarcinomas feature tumor cells with abundant intracellular mucin. The molecular mechanisms by which such mucin-producing lung adenocarcinomas develop remain unknown.

**Methods:** Using a Visium spatial transcriptomics approach, we analyzed IMA and compared it with SRCC patho-transcriptomically. Combining spatial transcriptomics data with *in vitro* studies using RNA-seq and ChIP-seq, we assessed downstream targets of transcription factors HNF4A and SPDEF that are highly expressed in IMA and/or SRCC.

**Results:** Spatial transcriptomics analysis indicated that there are 6 distinct cell clusters in IMA and SRCC. Notably, two clusters (C1 and C3) of mucinous tumor cells exist in both adenocarcinomas albeit at a different ratio. Importantly, a portion of genes (e.g., *NKX2-1*, *GKN1*, *HNF4A* and *FOXA3*) are distinctly expressed while some mucous-related genes (e.g., *SPDEF* and *FOXA2*) are expressed in both adenocarcinomas. We determined that HNF4A induces *MUC3A/B* and *TM4SF4* and that BI 6015, an HNF4A antagonist, suppressed the growth of IMA cells. Using mutant SPDEF that is associated with COVID-19, we also determined that an intact DNA-binding domain of SPDEF is required for SPDEF-mediated induction of mucin genes (*MUC5AC*, *MUC5B* and *AGR2*). Additionally, we found that XMU-MP-1, a SPDEF inhibitor, suppressed the growth of IMA cells.

**Conclusion:** These results revealed that IMA and SRCC contain heterogenous tumor cell types, some of which are targetable.

## Introduction

Lung adenocarcinoma, the leading cause of all cancer death (∼50,000 deaths/year in the US), is pathologically the most prevalent non-small cell lung cancer (60% of all lung cancer), followed by lung squamous cell carcinoma.^1,2^ Of lung adenocarcinoma, ∼5% is classified as invasive mucinous adenocarcinoma (IMA), which is characterized by tumor cells with abundant intracellular mucin.^3^ It may be difficult to histologically distinguish IMA from other mucin-producing lung adenocarcinomas, including adenocarcinoma with signet ring cell features (also called as signet ring cell carcinoma [SRCC] in the past)^4^ due to their similar cell type characteristics with abundant intracellular mucin.^5,6^ In the last decade, especially due to the advent of next-generation sequencing, driver oncogenes for IMA (e.g., mutant *KRAS*) have been revealed.^7,8^ The association of SRCC and various fusions, in particular *ALK* fusions have been reported^9–11^ and the gene expression profiles for IMA have been reported by our group and others.^5,12,13^ In general, the transcription factor NKX2-1 is absent in IMA while often present in SRCC.^5^ Beyond the gene expression pattern, the role of some of the transcription factors in IMA has been determined using mouse models. Loss of NKX2-1 in the presence of mutant *KRAS* drives IMA while the FOXA family and SPDEF cooperate with mutant *KRAS* to induce IMA.^13–17^ Identification of downstream gene targets of such transcription factors is required to understand the biology of IMA, which leads to potential therapeutics. Such transcription factors and potential downstream gene targets have been analyzed *in vitro* using cell lines and/or *in vivo* with immunohistochemistry; however, there is no guarantee that these downstream gene targets are indeed regulated by the transcription factors *in vivo* and *in situ* in humans. Notably, the recent development of spatial transcriptomics technology using the Visium spatial transcriptomics platform^18^ has enabled researchers to identify such downstream gene targets *in situ* on a genome-wide scale in different tissue types, including lung adenocarcinoma.^19–21^ This approach also helps us to understand the biology of IMA beyond experiments using mouse models and human cell lines. In the present study using the Visium spatial transcriptomics platform, we analyzed human IMA to identify which genes are expressed in IMA *in situ* compared to SRCC. We further validated downstream gene targets of the key IMA transcription factors HNF4A and SPDEF that were obtained by *in vitro* analysis using cell lines paired with *in situ* analysis of the spatial transcriptomics results to determine which genes can be modulated by potential therapeutics targeting HNF4A and SPDEF.

## Materials and Methods

### Human specimens

Paraffin and OCT (Optimum Cutting Temperature) sections of lung adenocarcinoma, including IMA and SRCC, were obtained from MGH (approval #2009P001838), University of Cincinnati Cancer Center Biospecimen Shared Resource (approval #TB0082), Hiroshima University (approval #E-1919) and Kawasaki Medical School (approval #1310) in accordance with institutional guidelines for use of human tissue for research purposes. Written informed consent was obtained from all participants. Patients’ information is summarized in Supplementary Table 1.

### Histology, immunohistochemistry and RNAScope analyses

Staining (H&E, Alcian blue and immunohistochemistry) was performed using 5 μm paraffin-embedded lung sections as described previously^14^ with antibodies for NKX2-1 (1:100, cat# 8G7G3-1, Seven Hills Bioreagents, Cincinnati, OH), GKN1 (1:100, cat# H00056287-M01, Abnova, Taipei City, Taiwan), plus additional antibodies for patient sample characterization: MUC5AC (1:1000 cat# ab3649, A00am, Waltham, MA), MUC5B (1:1000, cat# sc20119 Santa Cruz Biotechnology, Dallas, TX), FOXA3 (1:100 cat# sc-5361 Santa Cruz Biotechnology, Dallas, TX), HNF4A (1:100 EDTA, cat# PP-H1415-0C R&D Systems, Minneapolis, MN), FOXA2 (1:1000 cat# AF2400 R&D Systems, Minneapolis, MN), and ALK (1:100 EDTA, Cell Signaling Technologies, Danvers, MA). RNAScope for NKX2-1 mRNA (Advanced Cell Diagnostics [ACD], Newark, CA) was performed at University of Cincinnati Cancer Center Biospecimen Shared Resource (UCCC-BSR).

### Visium spatial transcriptomics analysis

Spatial gene expression was conducted using the 10x Visium Spatial Gene Expression Slide & Reagents Kit (cat# 1000187) for fresh frozen tissue according to the manufacturer’s directions. Briefly, frozen lung tumor tissue sections were cut at 10 µm thickness and placed on the Visium capture area within the 8 x 8 mm fiducial frames. Sections were fixed with methanol and stained with hematoxylin and eosin before brightfield images were scanned at 4X magnification. The tissue was permeabilized for 45 minutes based on 10x’s optimization protocol. cDNA synthesis, second strand synthesis, amplification (16 cycles), and library construction were done according to the kit protocol. Paired end sequencing was done on a NextSeq 500 sequencer at a depth of 150 million reads. Space Ranger (v1.3.0) from 10x Genomics was used to process and map each Visium sample. The pre-built human genome reference (refdata-gex-GRCh38-2020-A) from 10x Genomics was used for alignment and counting. Spots with more than 500 expressed genes and less than 20% of reads mapped to mitochondrial genes were included in the downstream analysis using Seurat (v4.2.0).^22^ Normalization of gene expression was performed on each Visium sample using Seurat SCTransform. Normalized gene expression profiles from different samples were integrated using the reciprocal principal component analysis (RPCA) based batch correction in Seurat. Spot clusters were identified in the integrated data using the Seurat FindClusters function with the SLM algorithm. Genes selectively expressed in each cluster were identified using the Seurat FindAllMarkers function using two-tailed Wilcoxon rank sum test with the following criteria: fold change >=1.2, Bonferroni adjusted p value <0.1, and expression percentage >20%. The DE test was performed for each cluster within each sample. Gene expression normalized by Seurat NormalizeData function was used for all DE tests. For the gene selectively expressed in each cluster, we considered the genes that passed the above DE criteria in both mucinous subtypes of lung adenocarcinoma, including invasive mucinous adenocarcinoma of the lung (IMA) and signet ring cell feature (SRCC). For each cluster, genes differentially expressed in IMA vs. SRCC were identified using Seurat FindMarkers function using two-tailed Wilcoxon rank sum test with the following criteria: fold change >=1.2, p value <0.05, and expression percentage >20%. Genes selectively expressed in the cluster in at least one Visium sample were included in the IMA vs. SRCC DE test for the cluster. Pseudo-time analysis of epithelial cell clusters (C0, C1, C3, and C5) was performed using Monocle3 (v1.0.0).^23^

### Cell culture, lentivirus infection, CRISPRi, TaqMan gene expression, siRNAs, immunoblotting, RNA-seq, ChIP-seq and ATAC-seq analyses

H2122 lung adenocarcinoma cells, A549 lung carcinoma cells and H292 lung mucoepidermoid carcinoma cells were obtained from ATCC (Manassas, VA). DV-90 lung adenocarcinoma cells were obtained from the DSMZ-German Collection of Microorganism and Cell Cultures GmbH (Science Campus Braunschweig-Süd, Germany). CRISPRi analysis was performed by transiently transfecting A549 cells that stably express dCas9-KRAB with custom TruGuide Synthetic gRNA (Thermo Fisher) designed by CRISPOR^24^ that targets the upstream region of the human *MUC5AC* gene where the transcription factor NKX2-1 binds (ChIP-seq datasets retrieved from GSE86959)^13,14^ using Lipofectamine^TM^ RNAiMAX Transfection Reagent according to the manufacturer’s instructions (cat# 13778075; Thermo Fisher Scientific, Waltham, MA). Forty-eight hours after transfection, RNA was extracted using TRIzol^TM^ Reagent (cat# 15596018; Thermo Fisher) and cDNA was made using iScript^TM^ cDNA Synthesis Kit (cat# 1708841; Bio-Rad, Hercules, CA). TaqMan gene expression analysis was performed using human gene probes, including *MUC5AC* (Assay ID # Hs00873651), *MUC5B* (Assay ID # Hs00861595), *SPDEF* (Assay ID # Hs00171942), *FOXA1* (Assay ID # Hs04187555), *FOXA2* (Assay ID # Hs00232764), *FOXA3* (Assay ID # 00270130) and *GAPDH* (Assay ID # Hs02758991) according to the manufacturer’s instructions (cat# 4453320 or 4331182, Thermo Fisher). Lentiviral vector carrying human HNF4A was made by inserting human *HNF4A8*^25^ (obtained from Addgene; cat# 31094) into the PGK-IRES-EGFP vector as described previously.^26^ Lentivirus was generated at the CCHMC viral vector core. H2122 cells were infected with lentivirus carrying *HNF4A* and treated with BI 6015 (cat# 4641, Tocris, Minneapolis, MN), Mycophenolic acid/MPA (cat# 1505, Tocris) or Sorafenib (cat# 6814, Tocris). Twenty-four hours after treatment, gene expression analysis was conducted as described above with additional TaqMan probes (cat# 4453320 or 4331182, Thermo Fisher), including *HNF4A* (Assay ID # Hs00230853), *MUC3A/B* (Assay ID # Hs03649367), *TM4SF4* (Assay ID # Hs00270335), *NKX2-1* (Assay ID # Hs00968940), *ACY3* (Assay ID # Hs00997729), *AGMAT* (Assay ID # Hs00383135), *METTL7B* (Assay ID # Hs00378551) and *CD274*/PD-L1 (Assay ID # Hs01125301). A549 cells were transfected with three different siRNAs targeting human *HNF4A* along with negative control siRNAs (cat# 4392420; siRNA ID: s6696, s6697 and s6698 for HNF4A and cat# 4390843 and 4390846 for negative control #1 and #2, respectively, Thermo Fisher) and gene expression analysis was performed as described above. A549 and DV-90 cells were also treated with BI 6015, mycophenolic acid/MPA or sorafenib as described above. Cell numbers were counted 6 days after cells were treated. Immunoblotting analysis with antibodies against HNF4A (cat# PP-H1415, R&D Systems, Minneapolis, MN), CDKN1A/p21 (cat# 2947, Cell Signaling Technology, Danvers, MA), Cleaved PARP (cat# 5625, Cell Signaling Technology) and ACTA1 (cat# A2066, MilliporeSigma, St. Louis, MO) was performed using extracts of cells treated for 24 hours. ChIP-seq was performed using antibodies against two different HNF4A antibodies (cat# PP-H1415 and AF4605, R&D Systems) and next-generation sequencing was performed at CCHMC DNA sequencing core (Cincinnati, OH) as described previously.^17^ Quality assessment and pre-processing of ChIP-seq reads were performed using FASTQC, Trim Galore and SAMtools. Reads were aligned to hg38 genome using Bowtie2. Low quality alignments and duplicate reads were removed using SAMtools and Picard MarkDuplicates tool. To identify HNF4A binding sites, peaks for each antibody were called using MACS2 (v2.1.0)^27^ callpeak function with the following parameters “-g hs -B --SPMR --keep-dup all -q 0.1”. ENCODE blacklist regions (accession number ENCFF356LFX) were filtered using bedtools intersect. Peaks with less than 2-fold enrichment and/or false discovery rate>=0.01 were also filtered, identifying 73,257 peaks bound by one HNF4A antibody (cat# PP-H1415) and 32,764 peaks by the other HNF4A antibody (cat# AF4605). Common peaks (n=29,426) bound by both antibodies were identified using Homer (v4.10)^28^ mergePeaks function with “-d given”. Motif enrichment was performed using Homer (v4.10) findMotifsGenome function. Visualization of ChIP-seq peaks and signals in genomic regions was performed using UCSC genome browser. ChIP-seq signal tracks were generated using MACS2 bdgcomp with “-m FE” to calculate fold enrichment in treatment over input control. Lentiviral vector carrying mutant SPDEF (G277D) was generated using QuikChange II XL Site-Directed Mutagenesis Kit (cat# 200521, Agilent, Santa Clara, CA) with oligonucleotides (5’-tcctcaattttgaagatgtccttctccttgttgagcc-3’ and 5’-ggctcaacaaggagaaggacatcttcaaaattgagga-3’ obtained from IDT, Coralville, Iowa) and wild type SPDEF cDNA.^29^ H292 cells were infected with lentivirus carrying wild type or mutant SPDEF (G277D) that was generated by the CCHMC viral vector core and RNA was made using extracted RNA as described above. RNA-seq was performed by CCHMC Genomics sequencing core as described previously.^13^ Quality assessment and pre-processing of RNA-seq reads were performed using FASTQC, Trim Galore and SAMtools. Reads were then aligned to hg38 genome using Bowtie2. Low-quality alignments were removed using SAMtools. Gene expression was counted using htseq-count. Differential expression analysis was performed using Bioconductor DESeq2 package.^30^ Differential expression with at least a 1.5-fold change and false discovery rate < 0.1 was considered significant. Heatmap visualizations of gene expression were performed using R package pHeatmap using variance stabilizing transformed expression values from the DESeq2 analysis. ATAC-seq was performed using ATAC-seq kit (cat# C01080002, Diagenode, Denville, NJ) with the CCHMC DNA sequencing core according to manufacturer’s instructions and analyzed as described previously.^31^ TaqMan gene expression analysis was conducted to validate some of the results from RNA-seq using probes mentioned above. A549 and DV-90 cells were treated with XMU-MP-1 (cat# 6482, Tocris) for 24 hours and RNA and protein were harvested. TaqMan gene expression and immunoblotting analyses were done with probes and antibodies mentioned above using extracted RNA and protein, respectively. Cell number was counted 6 days after treatment.

### Statistical analysis

Statistical differences were determined using two-tailed and unpaired Student’s or Welch’s *t*-test. Error bars represent SEM. The difference between two groups was considered significant when the *p*-value was < 0.05 (*). *P*-values of 0.001 to 0.01 are designated **, 0.0001 to 0.001 *** and < 0.0001 ****.

## Results

### Characteristics of IMA and SRCC

To understand the gene expression profiles of IMA and SRCC *in situ* on a genome-wide scale, we employed Visium spatial transcriptomics analysis^18^ using OCT embedded specimens of IMA and SRCC (Fig.1 and Supplementary Fig. 1). Bulk RNA-seq analysis indicated that the specimens used for Visium analysis carried a *KRAS* mutation (KRAS.Q61R) for IMA and *EML4-ALK* fusion for SRCC, respectively (Supplemental Table 1, top two samples). Alcian blue staining confirmed that both specimens contained abundant intracellular mucin (Fig. 1). As previously reported,^32,33^ the transcription factor NKX2-1 (also known as TTF-1) was negative in IMA tumor cells (of note, normal alveolar pneumocytes entrapped or interspersed between tumor cells are positive for NKX2-1; black nuclear staining in Fig.1) while NKX2-1 was positive in SRCC tumor cells (Fig. 1). Visium spatial transcriptomics analysis indicated that *ALK* mRNA was positive in SRCC but not in IMA (Fig. 1), which is consistent with bulk RNA-seq analysis (Supplemental Table 1) and in part validates the Visium analysis.

**Figure 1.**
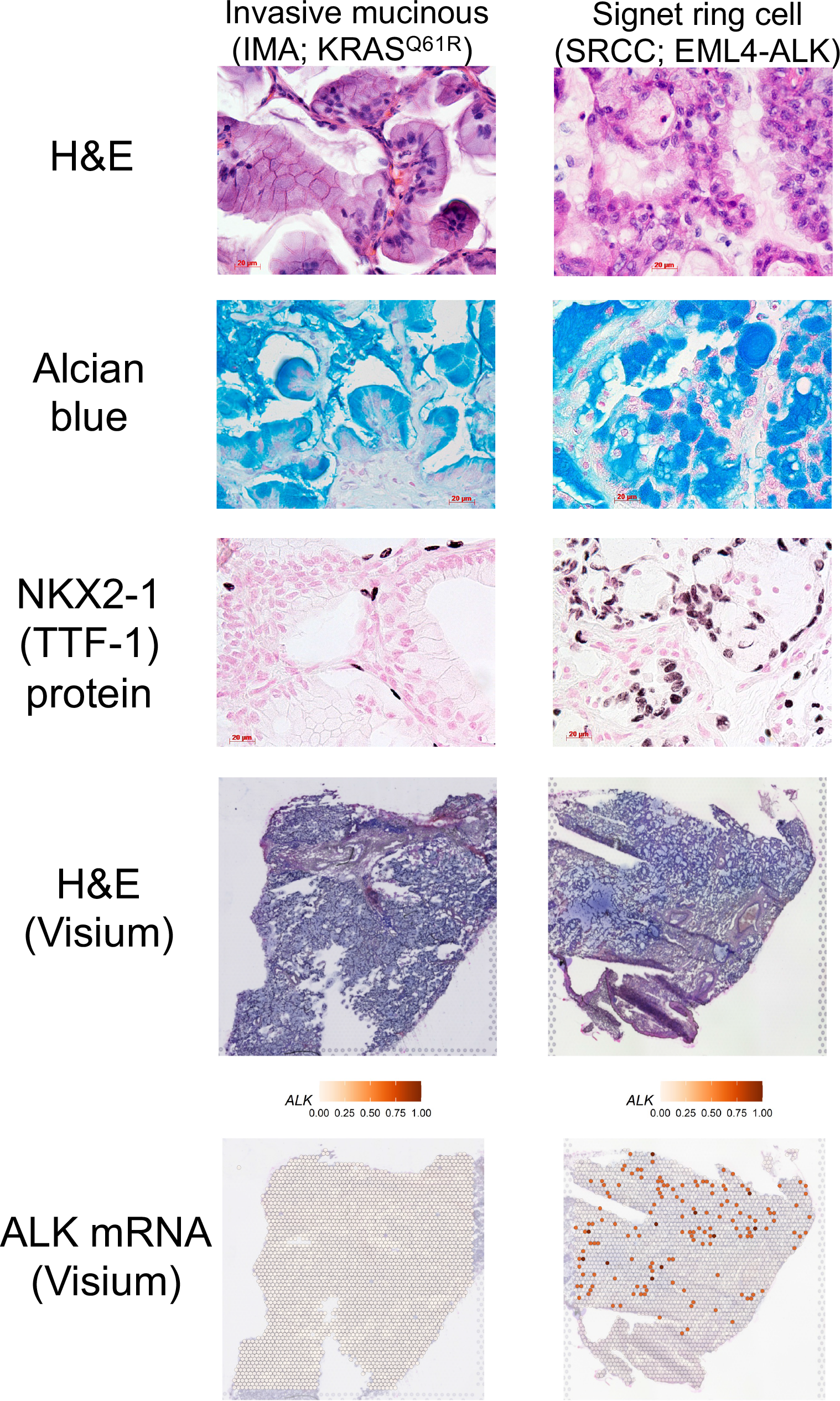
Similar but distinct pathology of invasive mucinous adenocarcinoma of the lung (IMA) and lung adenocarcinoma with signet ring cell features (SRCC) FFPE sections of IMA and SRCC were stained with H&E, Alcian blue and DAB with NKX2-1 antibody (top 3 panels). OCT sections of IMA and SRCC that were used for Visium spatial transcriptomics were visualized with H&E staining (second two panels from the bottom). The expression of *ALK* mRNA in SRCC but not in IMA was visualized using Seurat SpatialFeaturePlot (bottom panels).

### Visium based spatial transcriptomics of IMA and SRCC

We were able to obtain gene expression data from 5487 spots of IMA (2 sections) and 2572 spots of SRCC (1 section) at 55 μm resolution per spot (Fig. 2A-B and Supplementary Fig. 1). Unbiased clustering analysis identified 6 spot clusters, including tumor cells (C1 and C3), normal lung alveolar cells (C0), normal lung airway cells (C5), immune cells (C4) and stromal cells (C2), in IMA and SRCC. Unexpectedly, there were two tumor cell clusters (C1 and C3) in both IMA and SRCC regardless of their driver oncogene status (mutant *KRAS* vs. *ALK* fusion). Annotation of these clusters were validated by expression of cell type markers, including normal lung alveolar cells (C0 cluster; e.g., *SFTPC* and *AGER*), normal lung airway epithelial cells (C5 cluster; e.g., *SCGB1A1* and *FOXJ1*) (Fig. 2C, Supplementary Fig. 2), immune cells (C4; e.g., *PTPRC*/CD45 and *SPI1*/PU.1) and stromal cells (C2 cluster; e.g., *COL1A1* and *COL1A2*) (Fig. 2C, Supplementary Fig. 3). Differential expression analysis identified genes selectively expressed in each cluster in both IMA and SRCC (Supplementary Table 2) though most genes were expressed in different clusters at different degrees due to the limitation of Visium spatial transcriptomics at 55 μm resolution, a spot of which contains multiple cell types (Fig. 2D and Supplementary Table 2). Nevertheless, tumor cell populations of clusters C1 and C3 predominantly contained genes that were previously identified in IMA (e.g., *MUC5AC*, *MUC5B*, *MUC3A/B*, *HNF4A*, *FOXA3*, *SPDEF* and *FOXA2*) (Fig. 2C, D, Supplementary Fig. 4). With each cluster, we also identified genes differentially expressed between IMA and SRCC, e.g., in C3, *GKN1* expressed in IMA but not in SRCC (Supplementary Table 3). Trajectory analysis suggested that the normal lung alveolar cell population (C0) evolved into a tumor cell population (C3) through a transitory intermediate tumor cell population (C1) (C0 → C1 → C3) (Fig. 2E and Supplementary Table 2). These results indicate that mucin-producing lung adenocarcinomas (IMA and SRCC) harbor heterogenous cell populations, including two largely distinct tumor cell populations.

**Figure 2.**
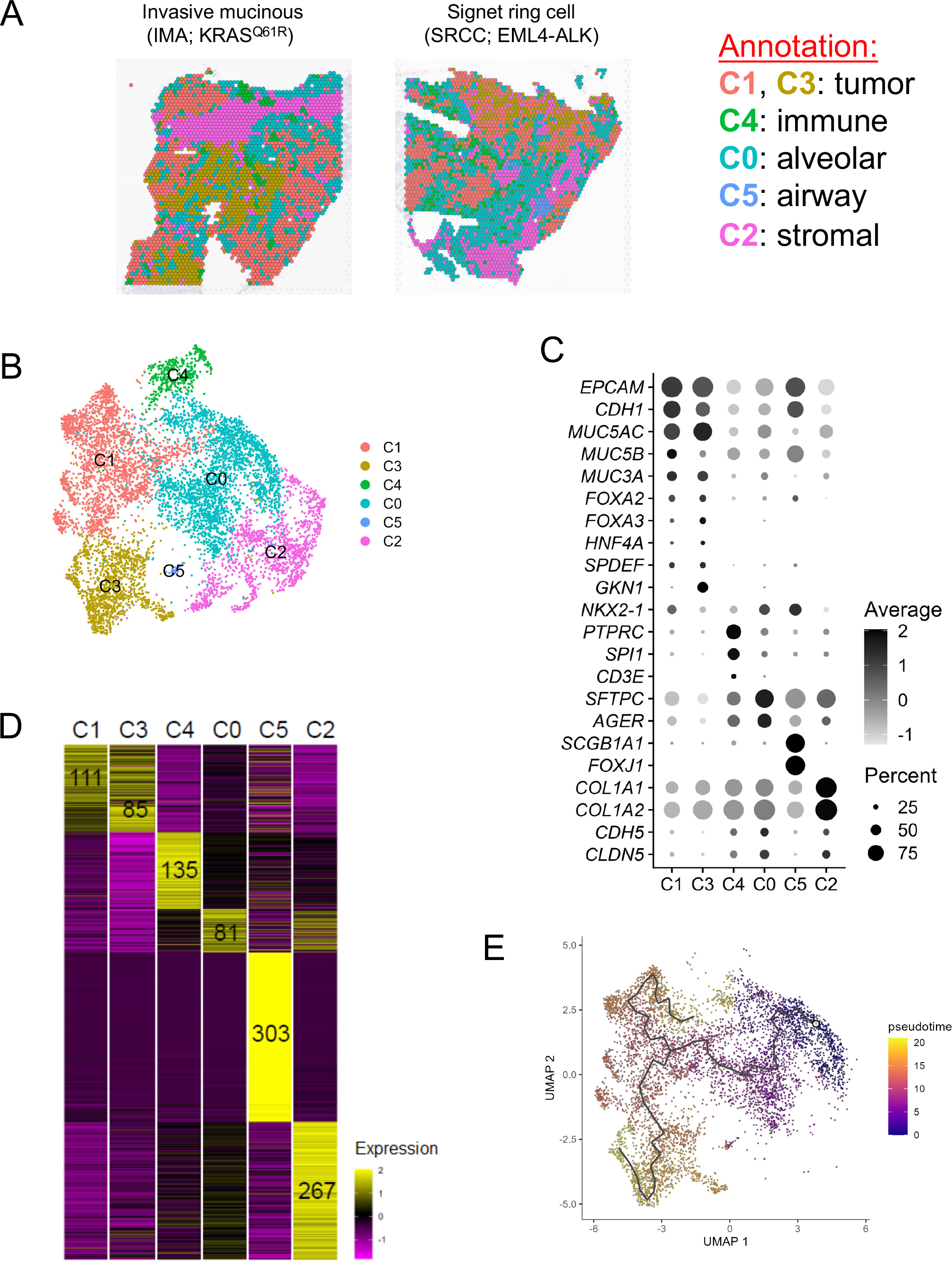
Spatial transcriptomics analysis of invasive mucinous adenocarcinoma of the lung (IMA) and lung adenocarcinoma with signet ring cell features (SRCC) A. Shown is clustering analysis of spatial transcriptomics data from OCT sections of IMA and SRCC (left panels). 6 spot clusters (C0 to C5) were identified and annotated (right panel). B. Shown is UMAP analysis of spatial transcriptomics data from OCT sections of IMA and SRCC. Cluster annotation is described in A. C. Average and percent of expression of representative genes in each cluster are shown. D. Heatmap visualization of scaled expression and numbers of genes selectively expressed in each cluster in both IMA and SRCC (padj<0.1, FC>=1.2, pct>=20%). E. Shown is pseudotime trajectory analysis of spatial transcriptomics data from OCT sections of IMA and SRCC. Normal alveolar cell cluster (C0) was used as a starting point.

### A subset of IMA expresses *NKX2-1* mRNA but not NKX2-1 protein

One of the most established pathogenetic characteristics of IMA is that NKX2-1 protein is absent in the majority of IMA tumor cells but not in SRCC tumor cells.^32,33^ In our Visium analysis, *NKX2-1* mRNA positive spots were reduced in IMA compared to those in SRCC (Fig. 3A and Supplementary Table 4), suggesting that loss of NKX2-1 protein in IMA occurs at the mRNA level.

**Figure 3.**
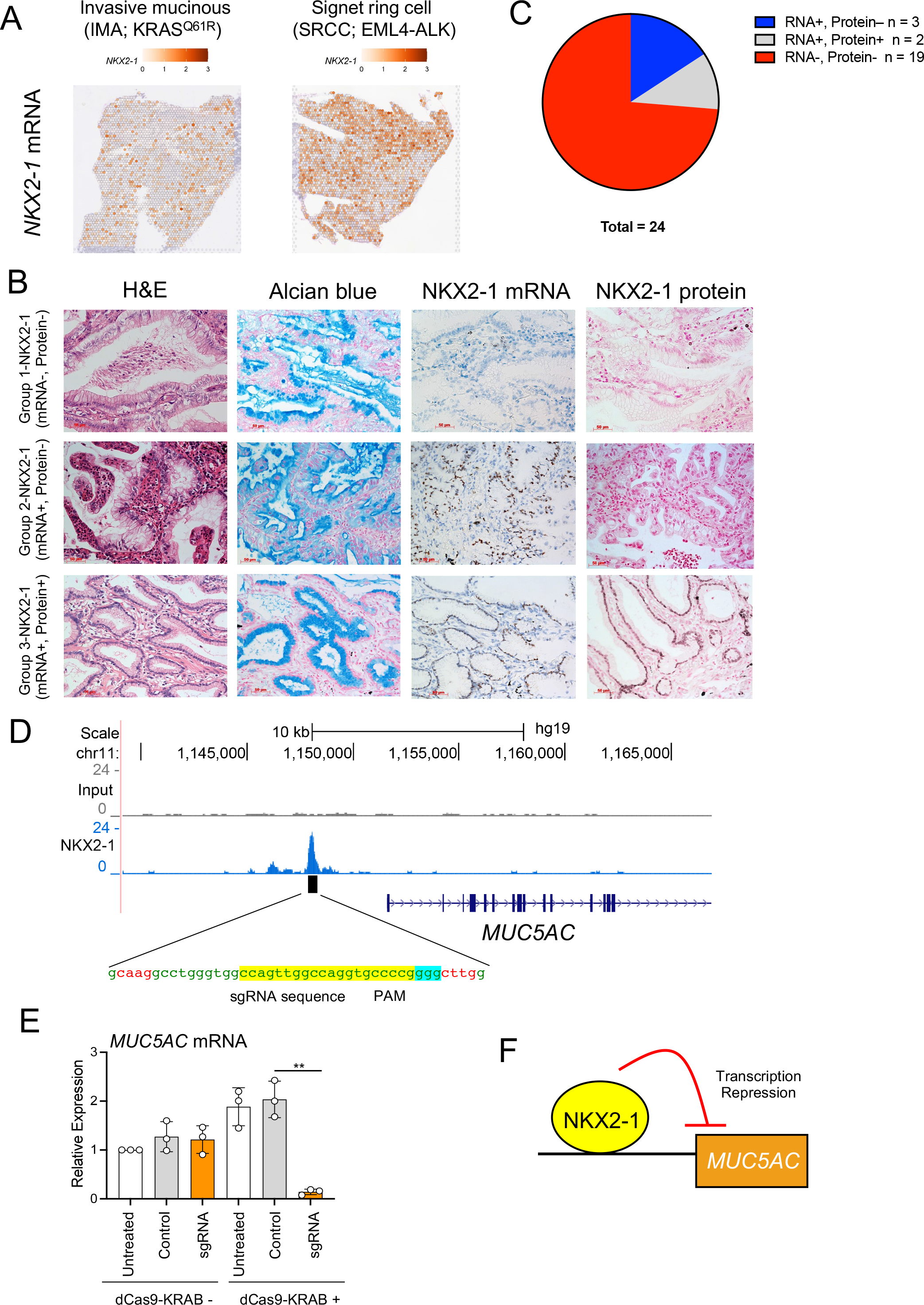
Expression of *NKX2-1* mRNA vs. protein in invasive mucinous adenocarcinoma of the lung (IMA) A. The expression of *NKX2-1* mRNA in IMA and SRCC is visualized using Seurat SpatialFeaturePlot. B. Three different groups of IMA specimens based on *NKX2-1* mRNA and NKX2-1 protein were analyzed with H&E and Alcian blue staining and *in situ* hybridization using RNAScope probe for *NKX2-1* mRNA and immunohistochemistry with NKX2-1 antibody. C. Shown is a pie chart describing the portion of three different groups of IMA specimens based on *NKX2-1* mRNA and NKX2-1 protein. ≥10% cutoff was used as positive expression. D. Shown is the sgRNA sequence targeting an NKX2-1 binding peak at the upstream region of *MUC5AC*, which was obtained by ChIP-seq analysis. E. Shown is the relative gene expression of *MUC5AC* from A549 human lung carcinoma cells transfected with the indicated synthetic sgRNA described in D with or without dCas9-KRAB (CRISPRi). F. Shown is a proposed model in which NKX2-1 represses the expression of *MUC5AC*.

However, recent reports indicate that a portion of IMA carry inactivating mutations in NKX2-1,^34,35^ making it possible that these IMAs express *NKX2-1* mRNA but not protein detectable by immunohistochemistry. To address this question, we looked at the expression of *NKX2-1* mRNA in IMA (FFPE specimens from 24 cases) *in situ* using RNAScope. Although the majority of IMA (79.2%) did not harbor *NKX2-1* mRNA (Fig. 3B-C and Supplementary Table 1), which is consistent with the Visium data (Fig. 3A), a portion of IMA mucin producing cells harbored *NKX2-1* mRNA but not NKX2-1 protein (12.5%) (Fig. 3B-C), suggesting that NKX2-1 loss in IMA occurs not only at the mRNA level but also at the protein level. Considering that IMA is also misdiagnosed as lung metastasis originating from non-lung cancers that do not carry *NKX2-1* (e.g., pancreatic cancer; https://www.proteinatlas.org/ENSG00000136352-NKX2-1/pathology),^36^ detection of *NKX2-1* mRNA using FFPE samples with RNAScope might be useful to distinguish primary IMA (*NKX2-1* mRNA positive) from lung metastasis (*NKX2-1* mRNA negative) in a sub portion of IMA.

Although the loss of NKX2-1 in the presence of mutant *KRAS* induces IMA as we reported previously,^14,15^ the mechanism by which NKX2-1 regulates genes involved in IMA is not fully understood. We have previously shown by ChIP-seq that NKX2-1 binds to the upstream non-coding region of *MUC5AC*, the most highly expressed gene in IMA^13^ (Fig.3D and Supplementary Fig. 5), suggesting that NKX2-1 represses the expression of *MUC5AC* by binding to this non-coding region. To assess whether this non-coding region affects the expression of *MUC5AC*, we employed a CRISPRi approach that we previously developed in which synthetic sgRNA targeting a non-coding gene regulatory region is transfected into dCas9-KRAB-stably expressing A549 lung carcinoma cells that express abundant *MUC5AC* and lack NKX2-1.^24^ Notably, the sgRNA targeting the NKX2-1 binding region upstream of *MUC5AC* significantly repressed the expression of *MUC5AC* (Fig. 3E and Supplementary Fig. 6), indicating this non-coding region functions as a gene-regulatory enhancer region to control the expression of *MUC5AC*. This result suggests that loss of NKX2-1 derepresses the expression of *MUC5AC* by liberating the enhancer region that was occupied by NKX2-1 (Fig. 3F).

### GKN1 is specifically expressed in a portion of IMA but not SRCC

Our clustering analysis using Visium data indicated that there are two clusters (C1 and C3) for tumor cell populations common in both IMA and SRCC (Fig. 2A-B). Nevertheless, cluster specific gene expression differences between IMA and SRCC can be identified (Supplementary Table S3). Among such genes, *GKN1* was specifically expressed only in the C3 cluster of IMA (Fig. 4A and Supplementary Table 3). *GKN1*, which is selectively expressed in normal human gastric mucus-secreting cells among all human normal cell types (Human Protein Atlas; see https://www.proteinatlas.org/ENSG00000169605-GKN1/single+cell+type), was previously shown to be expressed in a subset of mucinous tumor cells in an IMA mouse model and in 6 out of 11 human mucinous lung adenocarcinomas evaluated.^15^ In our Visium data, *GKN1* mRNA was expressed in IMA but not in SRCC (Fig. 4B). To further confirm this data, we looked at the expression of GKN1 protein by immunohistochemistry in 25 cases of IMA and 6 cases of SRCC that carry ALK fusions (Supplementary Table 1). Notably, GKN1 was positive in 10 out of 25 cases (40%) in IMA but negative in all cases of SRCC (Fig. 4C-D), suggesting, along with trajectory analysis (Fig. 2E), that mucinous tumor cells expressing GKN1 are the most well-differentiated mucinous tumor cell types among IMA.

**Figure 4.**
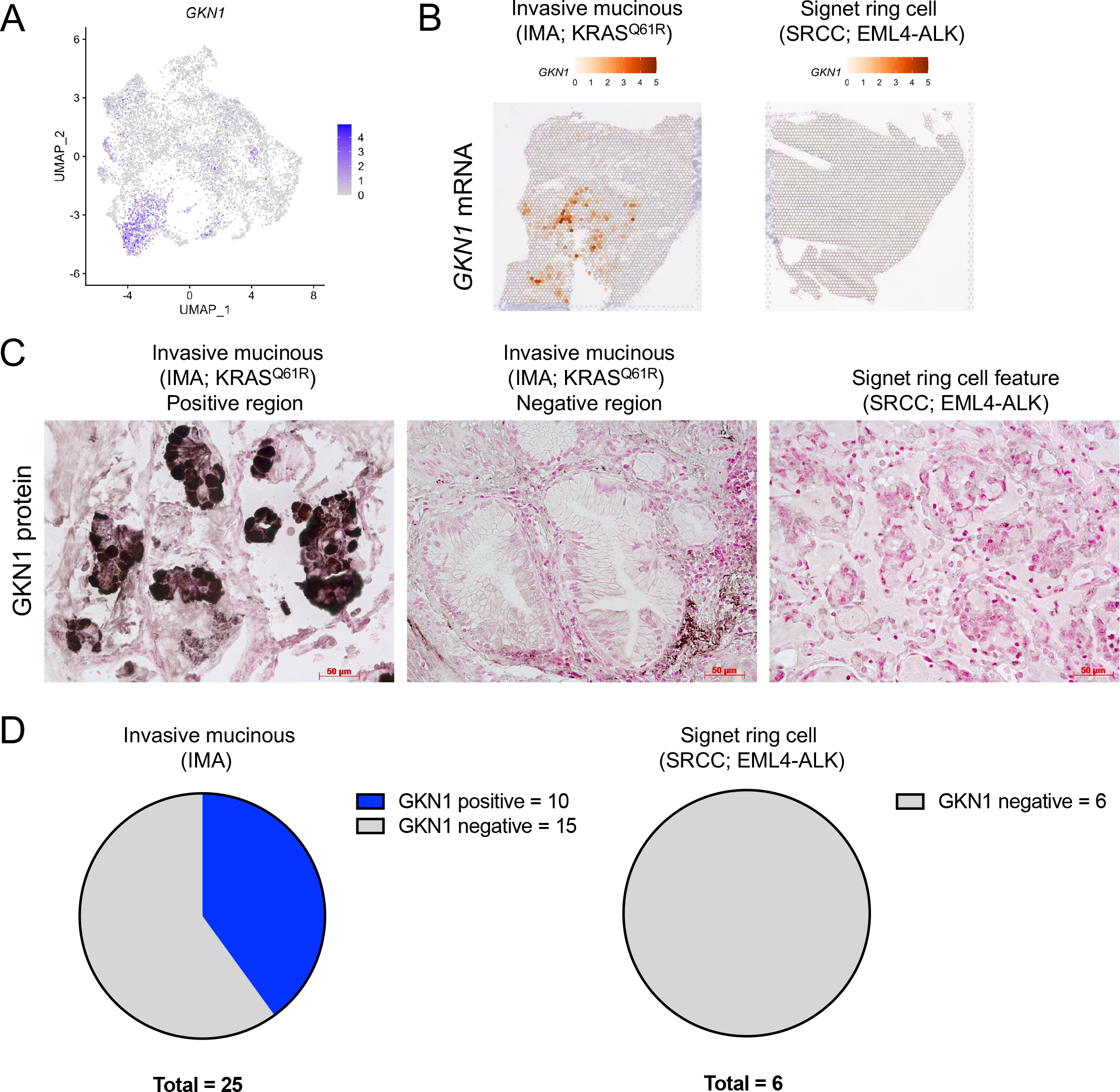
Selective expression of GKN1 in a portion of invasive mucinous adenocarcinoma of the lung (IMA) A. Shown is UMAP analysis of spatial transcriptomics data from OCT sections of IMA and SRCC, indicating that *GKN1*, a marker for gastric mucus-secreting cells, is selectively expressed in the C3 (but not C1) tumor cell cluster of IMA but not SRCC (see Supplementary Table 3). B. The expression of *GKN1* mRNA in IMA and but not SRCC is visualized using Seurat SpatialFeaturePlot. C. Shown is immunohistochemistry data indicating that GKN1 protein is expressed in a subset of IMA mucinous tumor cells but not all IMA mucinous tumor cells or SRCC mucinous tumor cells. D. Shown is a pie chart describing the number of GKN1 positive cases in IMA (10/25) and SRCC (0/6) patients.

### HNF4A induces *MUC3A/B* and *TM4SF4* in IMA

Expression of the transcription factor HNF4A, which is expressed in cells of normal gastrointestinal tract (GI), liver, pancreas and kidney but not in normal lung cells (Human Protein Atlas; https://www.proteinatlas.org/ENSG00000101076-HNF4A/tissue), is also one of the established pathogenetic characteristics of IMA but not of SRCC.^14,15,37^ Consistently, our Visium data analysis indicated that *HNF4A* mRNA was expressed in IMA but not in SRCC (Fig. 5A). Although lncRNA BC200 was reported as a critical downstream target of HNF4A in IMA using cell lines,^38^ its relevance *in situ* in IMA specimens is not yet understood. In order to understand the molecular mechanism by which HNF4A regulates genes that are involved in IMA pathogenesis *in situ* on a genome-wide scale, we induced the expression of HNF4A ectopically using lentivirus in H2122 lung adenocarcinoma cells that do not express endogenous HNF4A and determined the HNF4A downstream genes using RNA-seq (Fig. 5B and Supplementary Table 5). Then, we assessed their expression in our human IMA bulk RNA-seq data (6 cases)^13^ as well as the Visium platform. Among the genes induced by HNF4A in H2122 cells, more than 100 genes were highly expressed in IMA compared to normal lung. Among such IMA genes induced by HNF4A (Fig. 5B-C and Supplementary Fig. 7A-B), our Visium data indicated that *MUC3A/B* (expressed in normal GI but not in normal lung; Human protein atlas; https://www.proteinatlas.org/ENSG00000169894-MUC3A/tissue) and *TM4SF4* (expressed in normal GI and liver but not in normal lung; Human Protein Atlas; https://www.proteinatlas.org/ENSG00000169903-TM4SF4/tissue) were highly expressed in IMA but not in SRCC (Fig. 5A), both of which genes are reported to be involved in lung cancer progression.^39,40^ Our ChIP-seq data using H2122 cells that express ectopic HNF4A indicated that HNF4A directly bound to the promoter region of *MUC3A/B* and *TM4SF4* (Fig. 5D-E, Supplementary Fig. 7C-D and Supplementary Table 6), suggesting that HNF4A directly activates these genes and thereby promotes IMA tumorigenesis.

**Figure 5.**
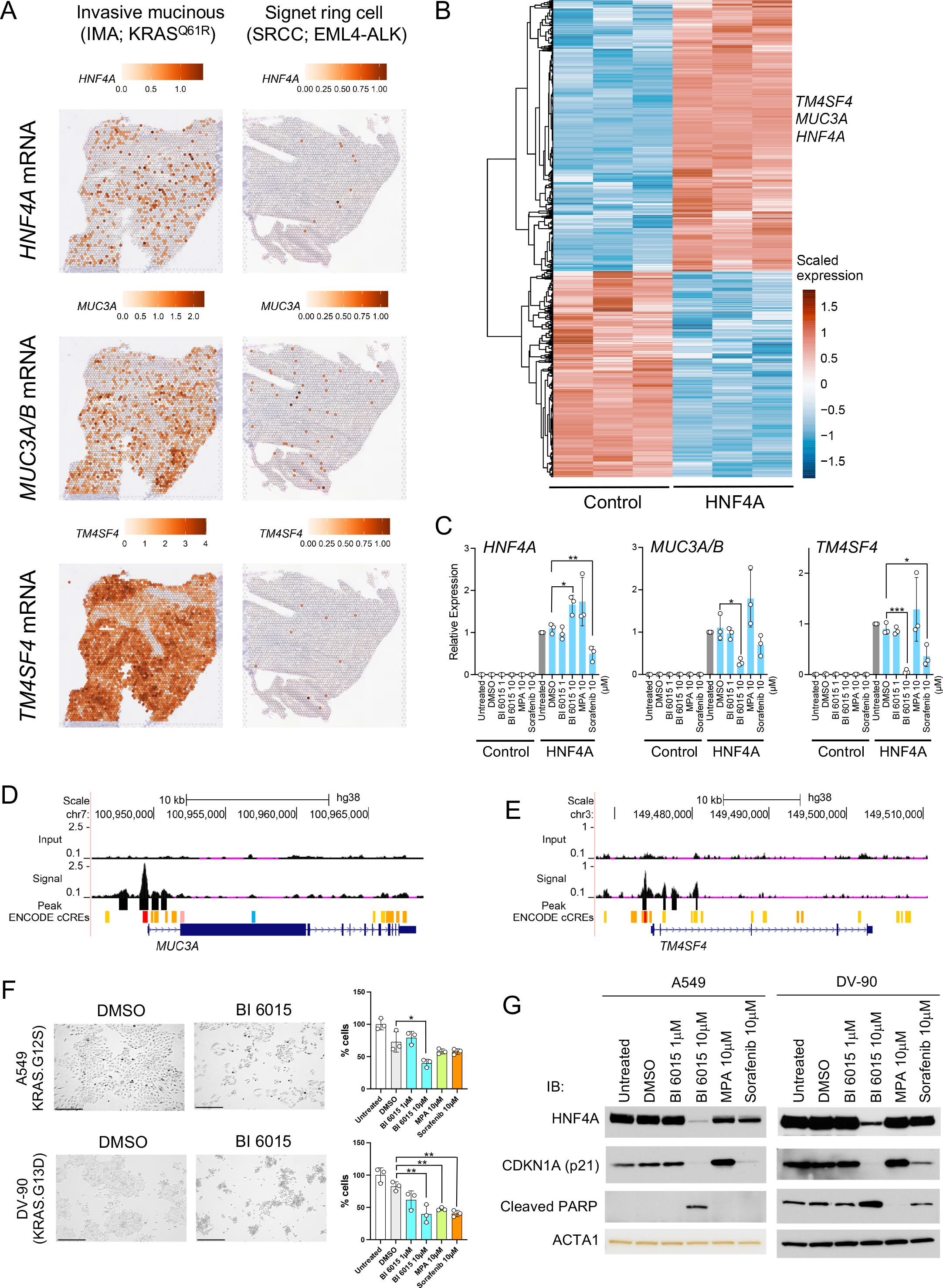
Induction of *MUC3A/B* and *TM4SF4* mRNA by HNF4A in IMA. A. The expression of *HNF4A*, *MUC3A/B* and *TM4SF4* mRNA in IMA and SRCC is visualized using Seurat SpatialFeaturePlot. B. Shown is RNA-seq data indicating differentially expressed genes (DEGs) that are regulated by HNF4A. RNA was isolated from H2122 lung adenocarcinoma cells (no endogenous HNF4A expression) that are infected with lentivirus carrying HNF4A or control lentivirus. DE was performed using DESeq2 and the DEG criteria were fold change >=1.5 and false discovery rate <0.1. C. Shown is TaqMan gene expression data indicating that *MUC3A/B* and *TM4SF4* mRNA were induced by HNF4A in H2122 cells as described in B in the presence of small molecule inhibitor/modifier compounds (BI 6015, MPA and sorafenib) targeting HNF4A (treated for 24 hours). D., E. Shown are ChIP-seq data indicating that HNF4A binds to the loci of *MUC3A/B* and *TM4SF4* in H2122 cells expressing HNF4A as describe in B. Two independent HNF4A antibodies were used for ChIP-seq (Shown was performed using PP-H1415 antibody; see Supplementary Fig. 7C for common regions bound by both antibodies). E. A549 and DV-90 lung adenocarcinoma cells that harbor KRAS mutations were treated with small molecule inhibitor/modifier compounds (BI 6015, MPA and sorafenib) targeting HNF4A and cells were counted. Representative images of the indicated cell lines that were treated with DMSO (control) or BI 6015 (10 µM) for 24 hours are shown (left panels). Cells were counted 6 days after treatment (right panels). The number of cells in the untreated group was set as 100%. Bar indicates 275 µM. F. Expression of endogenous HNF4A, CDKN1A (p21) and cleaved PARP was assessed by western blotting in A549 and DV-90 cells treated for 24 hours with small molecule inhibitor/modifier compounds (BI 6015, MPA and sorafenib) targeting HNF4A. ACTA1 was used as a loading control.

Multiple HNF4A antagonists, including BI 6015 and mycophenolic acid (hereafter, MPA) and a modulator, sorafenib, have been reported.^38,41,42^ We assessed whether these antagonists and the modulator affect HNF4A-mediated IMA gene expression, including that of *MUC3A/B* and *TM4SF4*. Notably, BI 6015 repressed the expression of both *MUC3A/B and TM4SF4,* plus other HNF4A downstream IMA genes *ACY3*, *AGMAT,* and *METTL7B* (Supplementary Fig. 7A-B), suggesting that BI 6015 is a gene-selective antagonist for HNF4A. The modulator, sorafenib, reduced HNF4A expression and downstream targets *TM4SF4, AGMAT,* and *METTL7B.* MPA did not repress any of the genes tested (Fig. 5C, Supplementary Fig. 7A and Supplementary Table 7). Since these antagonists and modulator have been reported to suppress tumor growth, including lung and liver cancer, we assessed whether they suppress IMA cell lines (A549 and DV90) that express endogenous HNF4A. Notably, all antagonists/modulator significantly reduced the growth of DV-90 cells while only BI 6015 significantly reduced the growth of A549 cells (Fig. 5F, Supplementary Fig. 8 and Supplementary Table 8). As reported for hepatocellular carcinoma cell lines,^41^ BI 6015 similarly reduced the expression of endogenous HNF4A and induced cleaved PARP, an apoptotic marker, in both IMA cell lines (A549 and DV-90 cells) (Fig. 5G and Supplementary Fig. 9-10), suggesting the HNF4A antagonist BI 6015 may be a potential therapeutic for IMA. Of note, another HNF4A antagonist MPA^38^ slightly reduced the expression of endogenous HNF4A and induced CDKN1A (p21), a cell cycle inhibitor, but not the apoptotic marker, cleaved PARP (Fig. 5G and Supplementary Fig. 9-10). These results suggest that HNF4A targeting antagonists can be potential therapeutics for IMA.

### An intact SPDEF DNA-binding domain is required for SPDEF-mediated induction of IMA genes

Previously, we determined that the transcription factor *SPDEF* is highly expressed in IMA and activates IMA genes (e.g., *MUC5AC* and *MUC5B*), thereby inducing IMA in the presence of mutant *KRAS*.^13^ Notably, our Visium spatial transcriptomics analysis indicated that *SPDEF* was expressed in both IMA and SRCC (Fig. 6A). While the pro-IMA transcription factor *FOXA2*^17^ was also expressed in both IMA and SRCC, another pro-IMA transcription factor *FOXA3*^13^ was highly expressed only in IMA (Fig. 6A). Importantly, *MUC5AC*, a major lung airway mucin,^13^ was highly expressed in IMA compared to SRCC while *MUC5B*, another major lung airway mucin, was highly expressed in SRCC but not in IMA (Fig. 6A). These results indicate similar but distinct pathogenesis of IMA and SRCC.

**Figure 6.**
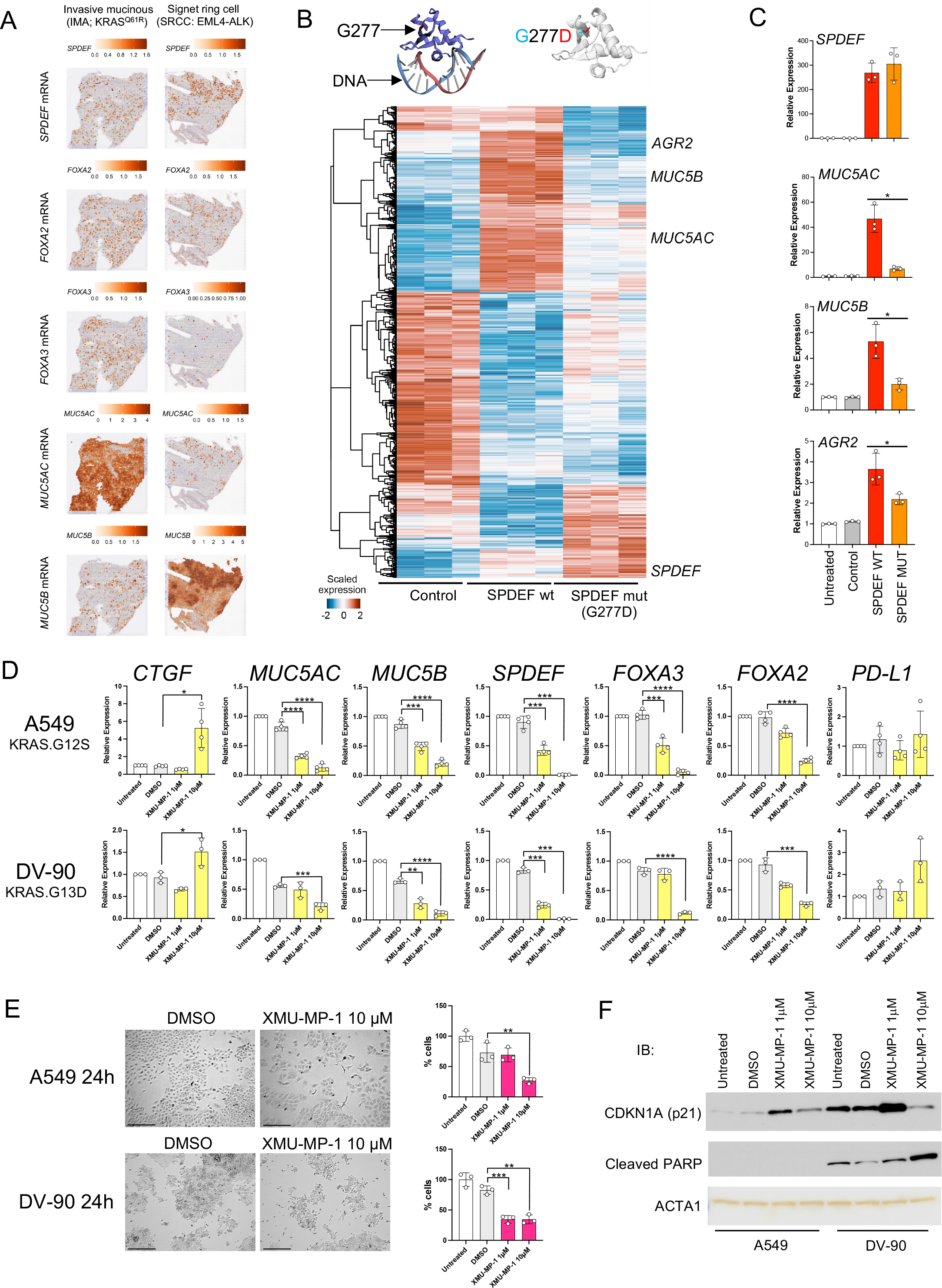
Mucous master regulator SPDEF as a potential therapeutic target for mucinous subtypes of lung adenocarcinoma. A. The expression of *SPDEF*, *FOXA2*, *FOXA3*, *MUC5AC* and *MUC5B* mRNA in IMA and SRCC is visualized using Seurat SpatialFeaturePlot. B. Shown is RNA-seq data indicating differentially expressed genes (DEGs) that are regulated by wild type SPDEF and/or DNA-binding domain mutant SPDEF (G277D). RNA was isolated from H292 lung mucoepidermoid carcinoma cells that were infected with lentivirus carrying wild type SPDEF or mutant SPDEF (G277D) or control lentivirus. DE was performed using DESeq2 and the DEG criteria were fold change >=1.5 and false discovery rate <0.1. C. Shown is TaqMan gene expression data indicating that *MUC5AC*, *MUC5B* and *AGR2* mRNA were highly induced by wild type SPDEF but not mutant SPDEF (G277D). D. Shown is TaqMan gene expression data indicating that the expression of mucous genes, *MUC5AC*, *MUC5B*, *SPDEF*, *FOXA3*, *FOXA2*, but not that of immune checkpoint ligand *CD274*/PD-L1 were significantly repressed by XMU-MP-1, an MST1/2 inhibitor (YAP/TAZ activator), in A549 and DV-90 lung adenocarcinoma cells treated for 24 hours. *CTGF*, a YAP/TAZ pathway downstream gene, was induced by XMU-MP-1 as expected. E. A549 and DV-90 lung adenocarcinoma cells that harbor *KRAS* mutations were treated with XMU-MP-1 as described in D and cell number was counted. Representative images of the indicated cells that were treated with DMSO (control) or XMU-MP-1 (10 µM) for 24 hours are shown (left panels). Cell number was counted 6 days after treatment (right panels). Counts of untreated groups were set as 100%. Bar indicates 275 µM. F. The expression of endogenous CDKN1A (p21) and cleaved PARP was assessed by western blotting in A549 and DV-90 cells treated for 24 hours with XMU-MP-1 (1µM, 10 µM). ACTA1 was used as a loading control.

Although SPDEF is a transcription factor, its non-nuclear function in innate immunity has been reported.^43^ Recently, a coding mutation that is associated with COVID-19 severity was reported in the DNA-binding domain of SPDEF (G277D),^44^ a possible structure of which is modeled in Fig. 6B, top panel, using AlphaFoldDB.^45^ Using this mutant, we assessed whether an intact DNA-binding domain of SPDEF is required for the induction of IMA genes. RNA-seq analysis using a H292 lung mucoepidermoid carcinoma cell line, a cell line in which endogenous *MUC5AC* induction by SPDEF is, to our knowledge most prominent,^29^ transduced with lentivirus carrying wild type SPDEF or mutant SPDEF (G277D) indicated that wild type, but not mutant SPDEF (G277D), induced the expression of IMA genes (*MUC5AC*, *MUC5B* and *AGR2*) (Fig. 6B, bottom panel, and Supplementary Tables 9-11) that is in part confirmed by TaqMan gene expression analysis (Fig. 6C and Supplementary Table 7), suggesting that the nuclear DNA-binding function of SPDEF is required for the induction of IMA genes. To assess whether SPDEF modifies chromatin accessibility to induce IMA genes, we also conducted ATAC-seq; however, chromatin accessibility was not influenced by SPDEF (Supplementary Fig. 11), suggesting that SPDEF binds to chromatin accessible regions but does not open chromatin as pioneer transcription factors such as the FOXA family do.

Recently, inhibition of the Hippo pathway MST1/2 kinase by the small molecule compound XMU-MP-1, thereby activating the transcriptional regulators YAP/TAZ, has been shown to repress the expression of *MUC5AC* and *SPDEF* in normal airway epithelial cells.^46^ We sought to determine whether XMU-MP-1 could also repress IMA genes in IMA cell lines (A549 and DV-90). Notably, XMU-MP-1 activated the expression of CTGF, a YAP/TAZ downstream gene, as expected, while it repressed the expression of not only *MUC5AC* and *SPDEF* but also other IMA genes *MUC5B*, *FOXA3* and *FOXA2* (Fig. 6D and Supplementary Table 7). Although IMA genes were repressed, an immune checkpoint inhibitor PD-L1, which is absent in IMA,^13^ was not induced (Fig. 6D and Supplementary Table 7). We further assessed whether XMU-MP-1 could suppress IMA tumor growth. Notably, XMU-MP-1 suppressed the growth of both A549 and DV-90 cells (Fig. 6E, Supplementary Fig. 12 and Supplementary Table 8), associated with the induction of CDKN1A (p21), a cell cycle inhibitor, in both cell lines and the cleavage of PARP, an apoptotic marker, in DV-90 cells but not in A549 cells (Fig. 6F and Supplementary Fig. 13). These results suggest that XMU-XP-1 can be a potential therapeutic for IMA.

## Discussion

IMA is pathologically well defined as a primary lung adenocarcinoma with tumor cells exhibiting goblet cell or columnar cell morphology and abundant intracytoplasmic mucin^3^; however, the molecular mechanisms, by which IMA develops, have just begun to be understood in the past decade. In thoracic pathology, the expression of NKX2-1 protein has been used to detect primary lung adenocarcinoma to distinguish lung metastasis while mucin-producing lung adenocarcinomas, in particular IMA, are often negative for NKX2-1 expression leading to difficult differentiation in diagnosis.^32^ A decade after the initial observation on the absence of NKX2-1 protein in IMA, we determined using human cell lines and mouse models that NKX2-1 is absent in IMA tumor cells because NKX2-1 functions as an anti-mucous transcription factor that represses the expression of mucous genes seen in IMA.^14,15^ Since this discovery, we have also determined that a number of transcription factors, including the FOXA family, SPDEF and HNF4A, induce the expression of IMA genes.^13,17^ However, only a limited number of downstream genes regulated by such transcription factors that were identified by RNA-seq using cell lines *in vitro* were validated pathologically, which hampers the effort to find potential therapeutics at a genome-wide scale for this specific type of lung adenocarcinoma. In the present study, using Visium spatial transcriptomics analysis, we determined the genes expressed in IMA and compared them with SRCC, which is a lung adenocarcinoma with abundant mucin production that may histologically mimic IMA, on a genome-wide scale, and linked them with the gene regulatory pathways that were identified by *in vitro* studies, which furthers our understanding of IMA and SRCC pathogenesis.

The recent technological development of spatial transcriptomics associated with next-generation sequencing, including the Visium platform,^18^ has provided gene expression profiles of different cell types in normal and diseased specimens, including lung cancer, on a sectioned slide *in situ*.^19–21^ Although the Visium platform used in this study was unable to identify cell types at a single cell level because of a resolution of only 55 μm,^18^ our analysis allowed us to locate different cell populations, including tumor cells, normal epithelial cells (airway and alveolar), immune cells and stromal cells, based on their gene expression in IMA and SRCC. This information helped us identify which non-tumor cell type in IMA (e.g., immune or stromal cell) expresses the IMA genes that we previously found using bulk RNA-seq (e.g., MMP9 from immune cells and PGF from stromal cells; Supplemental Table S3). Unexpectedly, the tumor cell populations of IMA and SRCC both contained the two clusters (C1 and C3) but carry distinct driver oncogenes (mutant *KRAS* and *ALK* fusion), suggesting that a shared tumorigenic pathway downstream of mutant *KRAS* and *ALK* fusion induces mucinous tumor cells. The trajectory analysis indicated a mucinous tumor evolutionary pathway of normal lung alveolar epithelial cells (C0) → tumor cells (C1) → tumor cells (C3). While IMA and SRCC contained both C1 and C3 spots, cluster selective genes specific in either IMA or SRCC can be identified. One of such genes was *GKN1*, which was expressed in the C3 cluster in IMA but not in other clusters in IMA or SRCC (Fig. 4). While GKN1 is downregulated in gastric cancer compared to normal gastric mucosa,^47^ the expression of GKN1 is associated with a poor prognosis in pancreatic cancer (Human Protein Atlas; https://www.proteinatlas.org/ENSG00000169605-GKN1/pathology). Further study to assess GKN1 as a prognostic factor for IMA and for its function in IMA tumorigenesis may be useful to delineate the role of the two different mucinous tumor cell clusters (C1 and C3) as to which cell cluster is associated with IMA and SRCC aggressiveness.

Mutant *KRAS* used to be an undruggable driver oncogene; however, the discovery of small molecule inhibitors for KRAS^G12C^ has revolutionized the treatment for non-small cell lung cancer (NSCLC) with mutant *KRAS*.^48^ Small molecular inhibitors targeting mutant KRAS, including KRAS^G12D^ (MRTX1133)^49^ that is frequently seen in IMA,^50^ are currently being tested in clinical trials (https://classic.clinicaltrials.gov/ct2/show/NCT05737706) and expected to be clinically available in the near future. Thus, these small molecular inhibitors targeting mutant KRAS (sotorasib, adagrasib or MRTX1133), ALK fusions (e.g., lorlatinib) or other drivers are becoming a first line therapy for unresectable NSCLC regardless of histologic subtypes. However, tumors recur in most cases and conventional chemotherapy is the only option when the molecularly-targeted therapy is not available. In the present study, we sought to identify small molecule inhibitors for the transcription factors HNF4A and SPDEF that are highly expressed in IMA. These small molecule inhibitors repressed their downstream target genes and suppressed the growth of IMA cell lines, which warrants these drugs as potential therapeutics for IMA along with driver oncogene-targeting molecular approaches especially for recurring IMA. Although XMU-MP-1 (a SPDEF inhibitor) does not appear to be a strong apoptosis inducer, it reduced the expression of mucous genes such as *MUC5AC* and *MUC5B*, which may alter the tumor microenvironment and enhance the effect of molecularly-targeted drugs and immune checkpoint inhibitors.

SPDEF with its ETS DNA binding domain is the transcription factor that is required for mucous gene expression.^29^ We have previously shown that SPDEF binds to the loci of two major mucin genes *MUC5AC* and *MUC5B* and induces their expression.^13^ The SPDEF ETS DNA binding domain is assumed to be required for such mucous gene expression; however, due to its non-nuclear function affecting the innate immunity pathway through MYD88,^43^ the role of the ETS DNA binding domain on mucus gene regulation remains uncertain. Recently, a mutation in the ETS DNA binding domain (G277D) was identified in patients who are severely affected by COVID-19.^44^ In the present study using SPDEF that carries the mutation, we sought to determine whether an intact ETS DNA binding domain is required for mucous gene expression. While wild type SPDEF induced the expression of the mucous genes, the mutant SPDEF did not, indicating that an intact ETS DNA binding domain is required for mucous gene expression. These results also suggest that the population which carries SPDEF^G277D^ may have a defect to induce mucous genes properly in response to respiratory diseases, including COVID-19, thereby resulting in severe symptoms while the mutation might have a protective effect against chronic lung diseases associated with hyper-mucus secretion, including asthma, COPD, IPF and mucin-producing lung adenocarcinomas such as IMA and SRCC, due to lower expression of mucous genes caused by the absence of functional SPDEF.

The continuing development of new technologies associated with next-generation sequencing (e.g., single cell-seq and spatial transcriptomics), CRISPR/Cas9 genome editing, and mouse models has enabled us to understand the biology of IMA, pathogenetically beyond pathological analysis and *in vitro* studies using cell lines. Although the current major treatment for unresectable IMA is conventional chemotherapy, the present study also provides potential therapeutics targeting highly expressed genes such as HNF4A and SPDEF. The current datasets have also provided tumor-associated cell populations such as immune and stromal populations in addition to tumor cell populations. Along with genetically-engineered mouse models that recapitulate the tumor-associated environment, studies that target these cell populations may lead to novel therapeutics such as immunotherapy for IMA beyond molecularly-targeted therapy.

## Author contributions

Y. Maeda, M. Mino-Kenudson, E.L. Snyder, T. Tsuchiya and T. Fukazawa designed experiments. W.D. Stuart, I.M. Fink-Baldauf and Y. Maeda performed experiments. M. Guo and H. Watanabe performed bioinformatical analyses. Y. Maeda, W.D. Stuart, I.M. Fink-Baldauf analyzed data. M. Ito, M. Okada, M. Mino-Kenudson and T. Fukazawa provided human specimens. All authors contributed experimental interpretation and manuscript writing.

## Acknowledgments

This work was supported by NIH grants (U01HL134745, R01CA240317), University of Cincinnati Cancer Center Pilot Project Award Program grant (S323349-2023 Ride Cincinnati-Maeda), Cincinnati Children’s Hospital Medical Center (Trustee Award Grant, CF-RDP Pilot & Feasibility Grant, GAP funding) to Y. Maeda. This research was made possible, in part, using the Cincinnati Children’s Genomics Sequencing Core RRID# SCR_022630, Viral Vector Core SCR_022641, Bio-Imaging and Analysis Facility SCR_022628, and the Integrated Pathology Research Facility SCR_022637. We thank Matt Kofron, Rafael Rosell, Betsy DiPasquale, Xiang Ao and Mary Durbin for assistance and discussions.

## Notes

### Competing Interest Statement

The authors have declared no competing interest.

