## Supplementary Figures for "Patho-transcriptomic analysis of invasive mucinous adenocarcinoma of the lung (IMA): comparison with lung adenocarcinoma with signet ring cell features (SRCC)"

Stuart et al., 2024

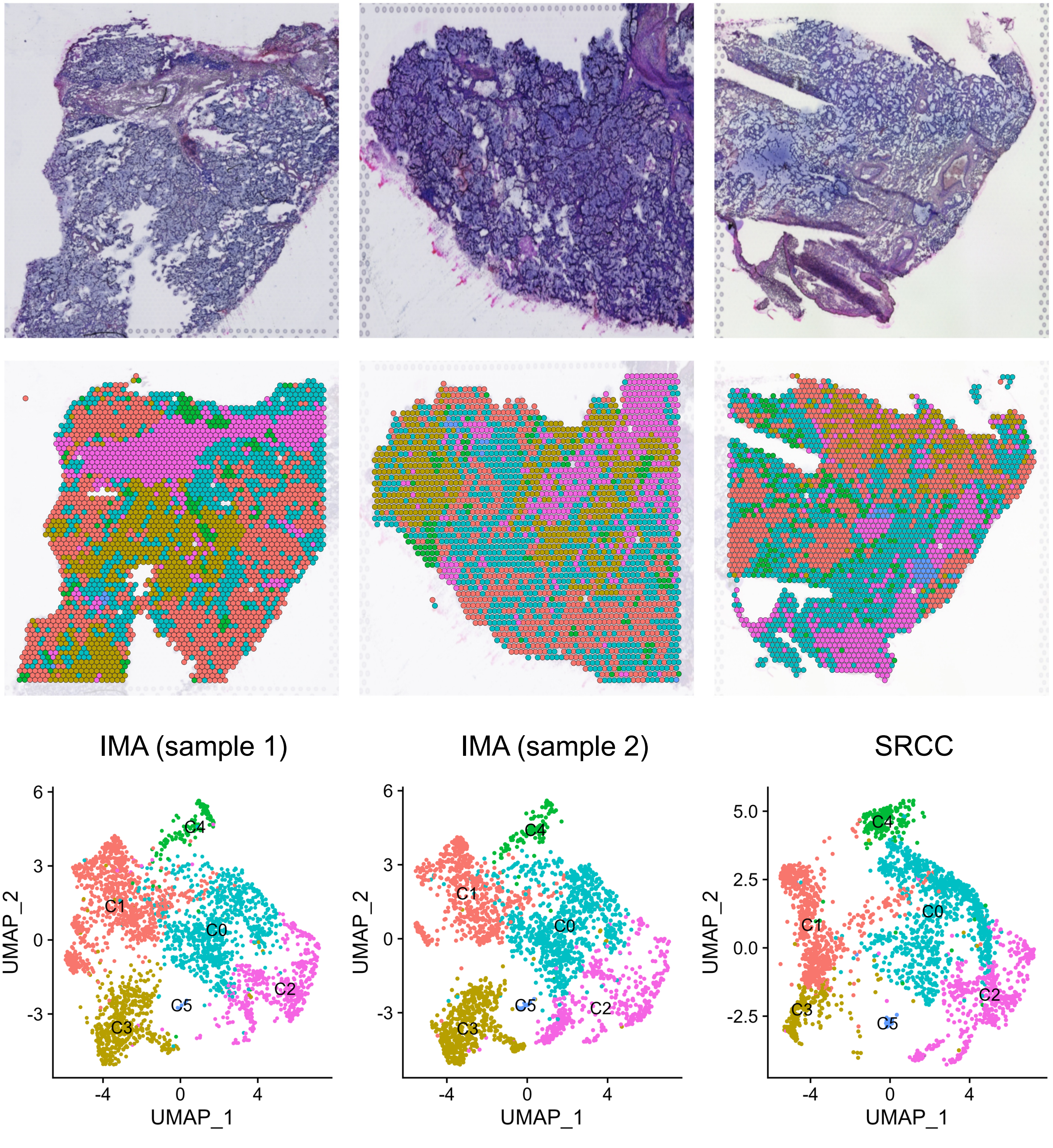


**Supplementary Figure 1. UMAP of spatial transcriptomics data from OCT sections of invasive mucinous adenocarcinoma of the lung (IMA) and lung adenocarcinoma with signet ring cell features (SRCC) indicates 6 different cell clusters (C0 to C5).**
Two left columns show data of two different sections from an IMA patient and the right column shows data of one section from an SRCC patient.

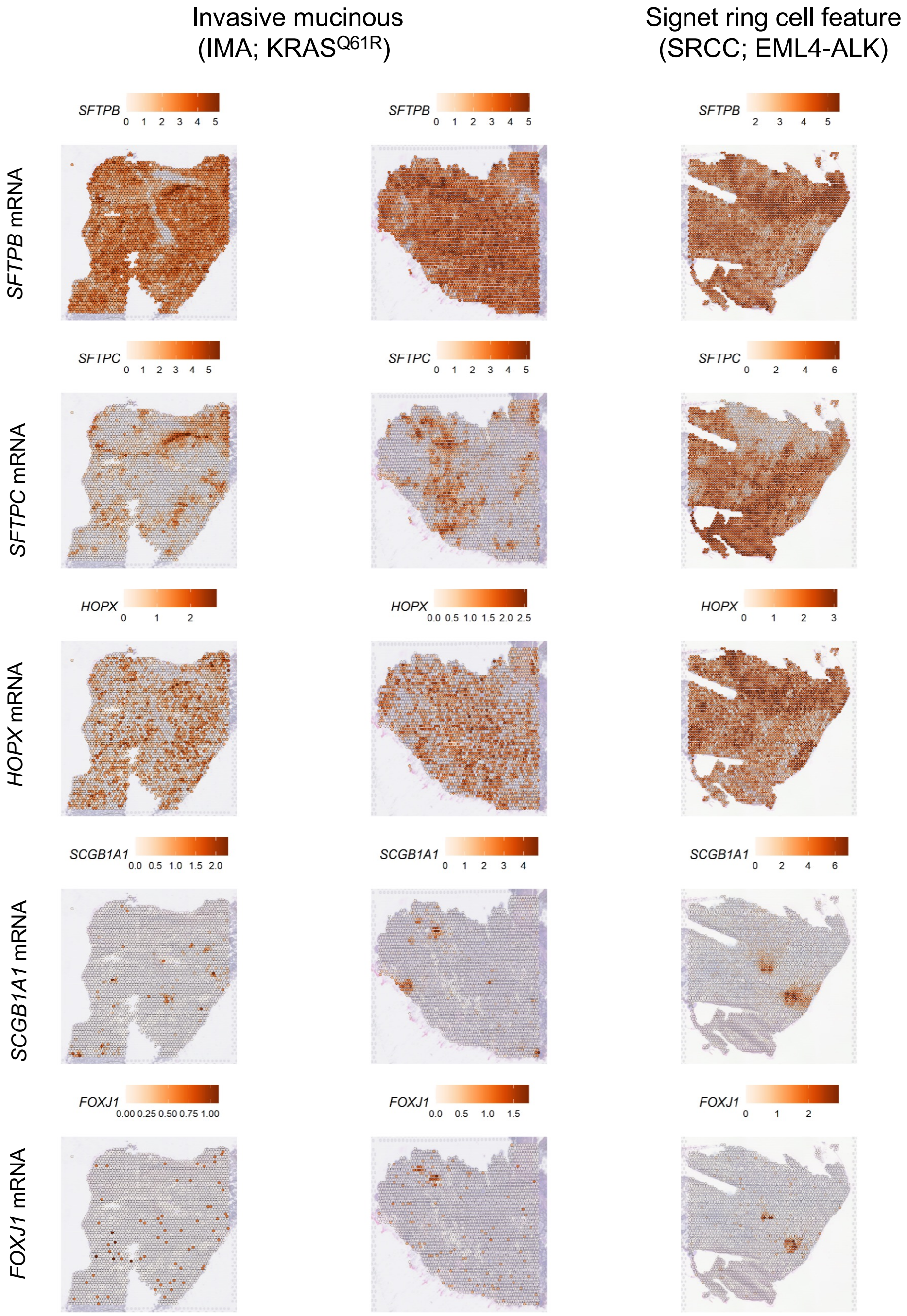


**Supplementary Figure 2. The expression of normal epithelial cell genes' mRNA in the spatial transcriptomics of IMA and SRCC.**
Two left columns show data of two different sections from an IMA patient and the right column shows data of one section from an SRCC patient. *SFTPB* is a marker for alveolar epithelial cells. *SFTPC* is a marker for alveolar type 2 cells. *HOPX* is a marker for alveolar type 1 cells. *FOXJ1* is a marker for airway ciliated cells.

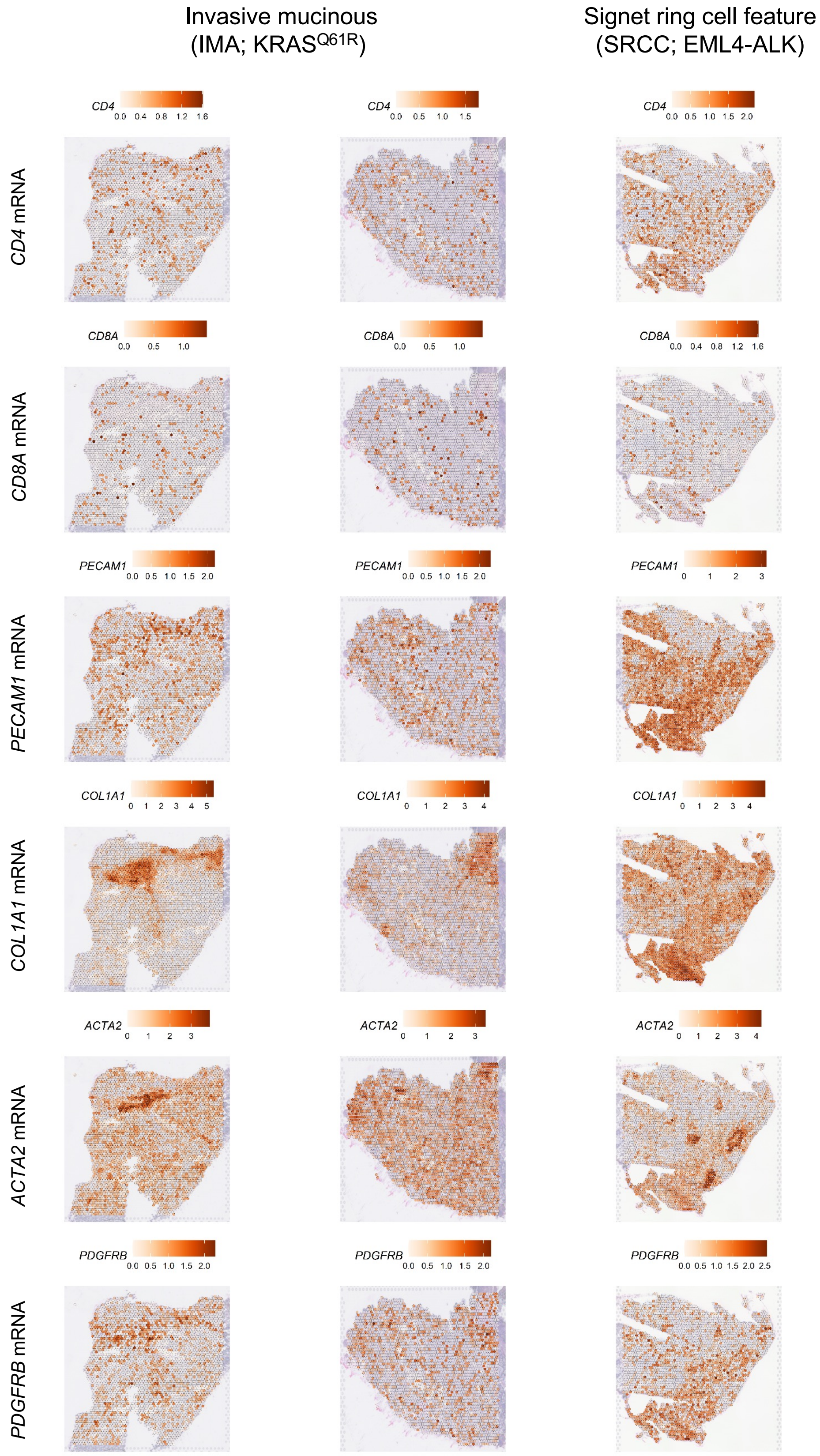


**Supplementary Figure 3. The expression of non-tumor cell genes' mRNA in the spatial transcriptomics of IMA and SRCC.**
Two left columns show data of two different sections from an IMA patient and the right column shows data of one section from an SRCC patient. *CD4* and *CD8* are markers for T cells. *PECAM1* is a marker for endothelial cells. *COL1A1*, *ACTA2* and *PDGFRB* are markers for mesenchymal cells, including fibroblasts and smooth muscle cells.

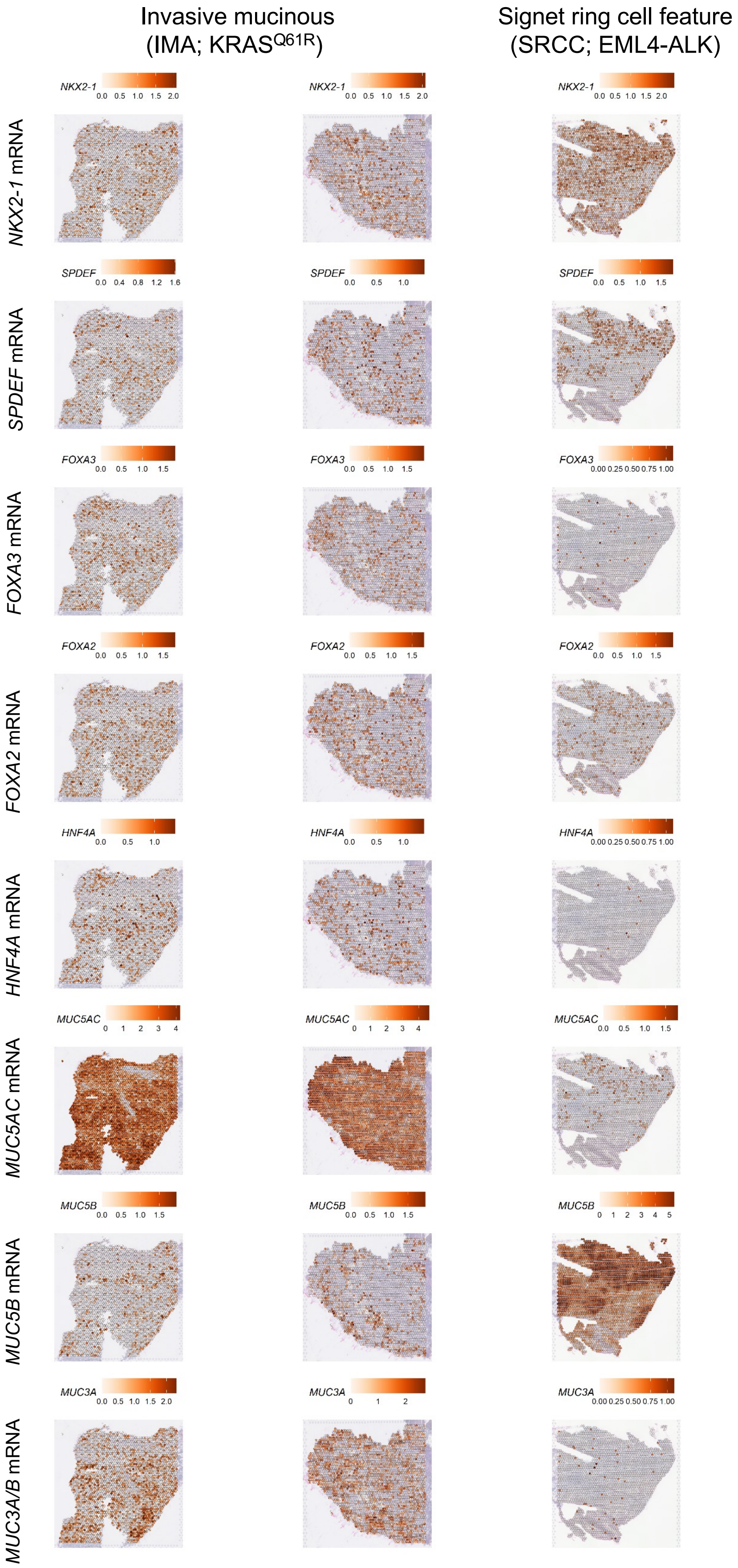


**Supplementary Figure 4. The expression of mucinous tumor suppressive/tumorigenic genes' mRNA in the spatial transcriptomics of IMA and SRCC.**

Two left columns show data of two different sections from an IMA patient and the right column shows data of one section from an SRCC patient. *SFTPB* is a marker for alveolar epithelial cells. *SFTPC* is a marker for alveolar type 2 cells. *HOPX* is a marker for alveolar type 1 cells. *FOXJ1* is a marker for airway ciliated cells.

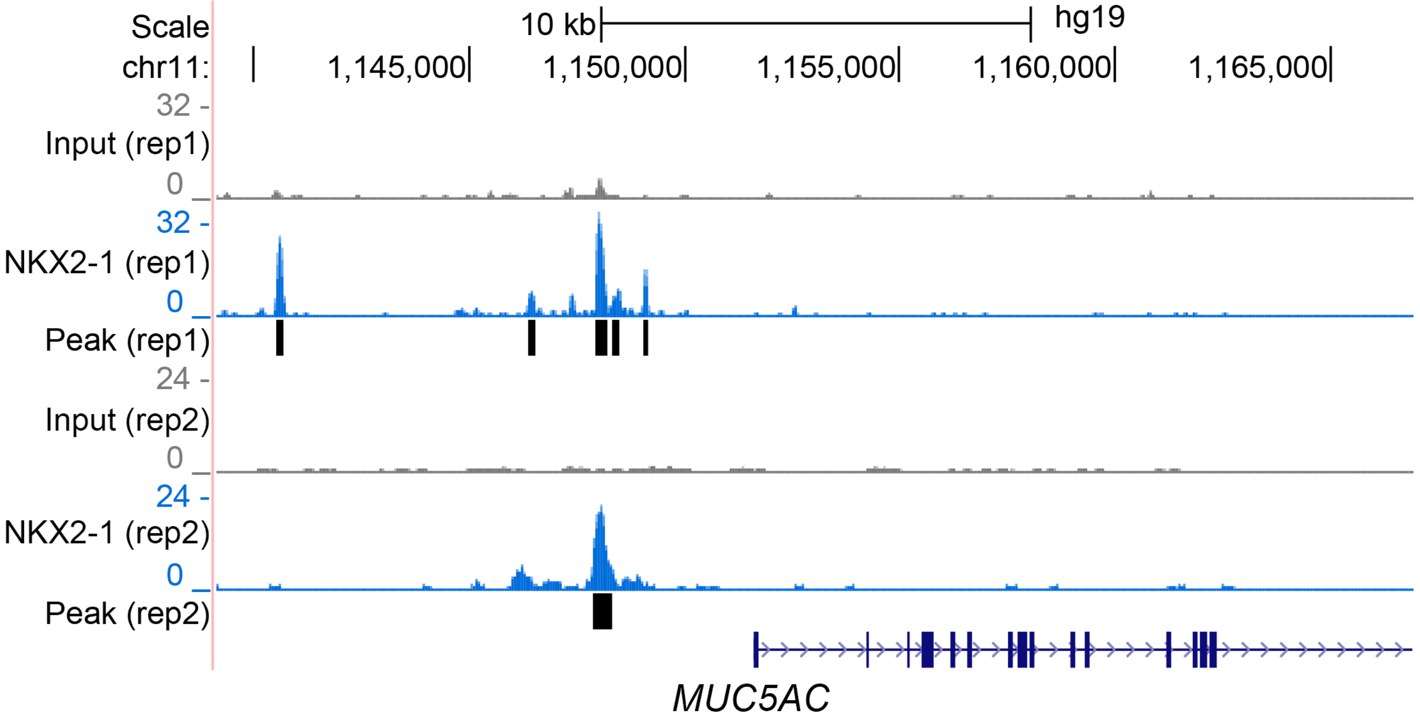


**Supplementary Figure 5. NKX2-1 binds to the upstream region of *MUC5AC*.**
Shown is a ChIP-seq peak indicating that NKX2-1 binds to the 3 kb upstream region of *MUC5AC*. ChIP-seq data was obtained from our previous ChIP-seq analysis[13, 14] that was performed using A549 lung carcinoma cells that stably express ectopic NKX2-1.

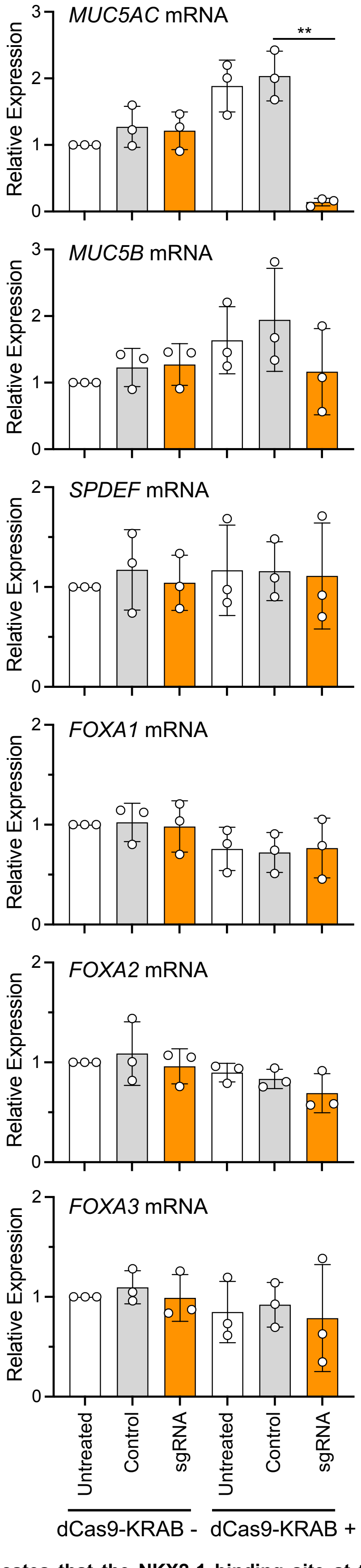


**Supplementary Figure 6. CRISPRi analysis indicates that the NKX2-1 binding site at the upstream region of *MUC5AC* is an enhancer region.**

Shown is TaqMan gene expression data that indicates sgRNA targeting the NKX2-1 binding site represses the expression of only *MUC5AC* but not other genes, including *MUC5B*, *SPDEF*, *FOXA1*, *FOXA2* or *FOXA3*, in A549 lung carcinoma cells that stably express dCas9-KRAB, suggesting NKX2-1 represses the expression of *MUC5AC* directly binding to the enhancer region of *MUC5AC* but not through the repression of pro-mucous transcription factors. *GAPDH* mRNA was used for normalization. Control indicates non-targeted sgRNA.

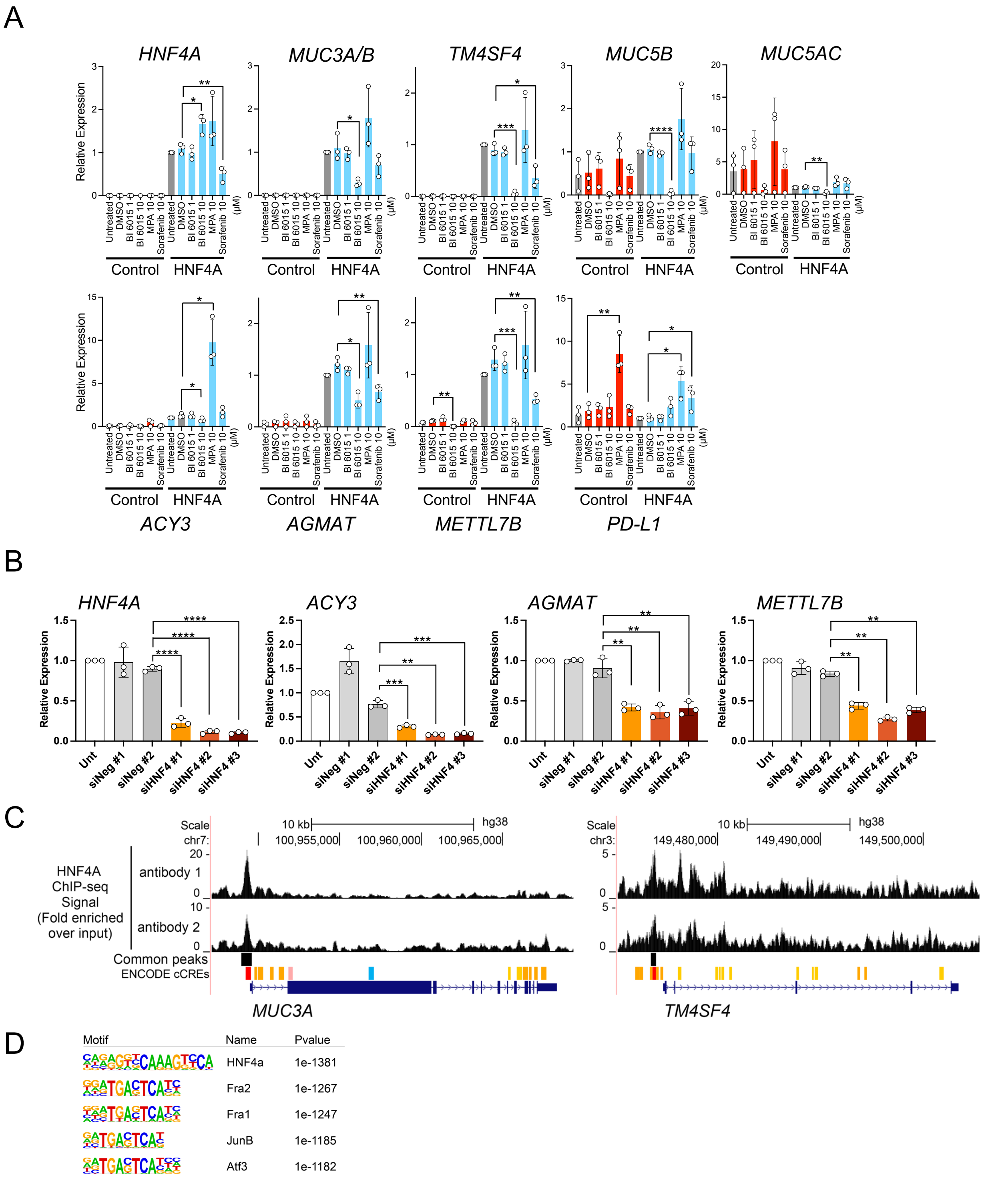


**Supplementary Figure 7. Nuclear receptor HNF4A induces the expression of IMA genes.**

A. Shown is TaqMan gene expression data indicating that HNF4A induces the expression of genes in H2122 lung adenocarcinoma cells that are expressed in invasive mucinous adenocarcinoma of the lung (IMA), which was confirmed by bulk RNA-seq using IMA specimens and Visium spatial transcriptomics analysis. H2122 cells were treated with small molecule inhibitor/modifier compounds (BI 6015, MPA and sorafenib) targeting HNF4A for 24h and the effect on expression of the indicated genes was determined. *18S* mRNA was used for normalization.

B. Shown is TaqMan gene expression data indicating that 3 independent siRNA-mediated knockdowns of *HNF4A* repressed the expression of IMA genes, including *ACY3*, *AGMAT* and *METTL7B*, in A549 lung carcinoma cells that express endogenous *HNF4A*. *GAPDH* mRNA was used for normalization.

C. Shown are ChIP-seq data indicating that HNF4A binds to the loci of *MUC3A/B* and *TM4SF4* in H2122 cells expressing HNF4A as described in Fig. 5B. The signal tracks represent fold of enrichment of signal in treatment over input control. Two independent HNF4A antibodies (antibody 1: PP-H1415 and antibody 2: AF4605) were used for ChIP-seq. Binding regions by each antibody were identified using MACS2 and common binding regions were identified using Homer mergePeaks.

D. Shown are top 5 transcription factor binding motifs most enriched by the common binding regions obtained from ChIP-seq data in which genome-wide HNF4A8 binding sites were assessed in H2122 cells using two independent antibodies.

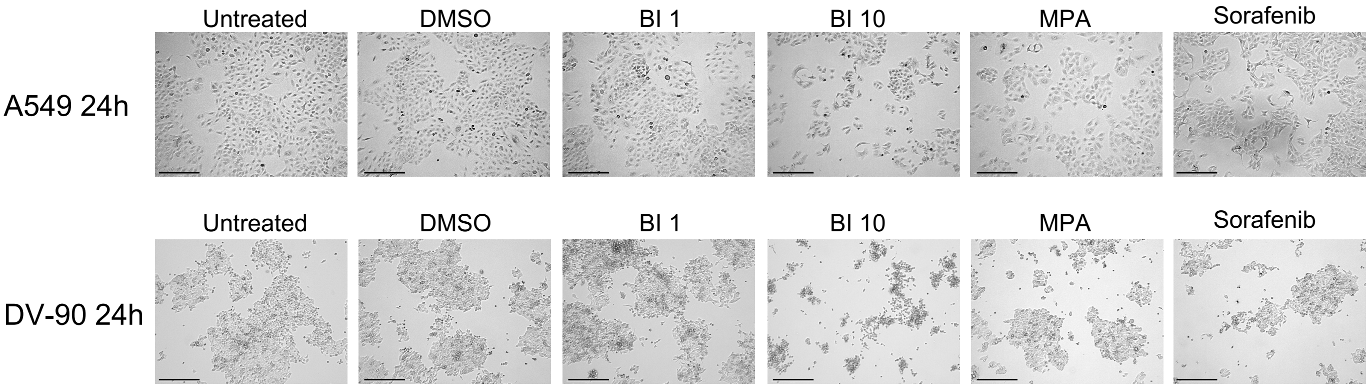


**Supplementary Figure 8. Representative images of A549 lung carcinoma cells and DV-90 lung adenocarcinoma cells treated with HNF4A-targeting small molecule inhibitor/modifier compounds.**

A549 and DV-90 cells that harbor KRAS mutations were treated with small molecule inhibitor/modifier compounds (BI 6015, MPA and sorafenib) targeting HNF4A. Shown are representative images of the indicated cells that were treated with DMSO (control), BI 6015 (1 μM or 10 μM), MPA (10 μM) or sorafenib (10 μM) for 24 hours. Bar indicates 275 μM.

1st independent experiment

2nd independent experiment

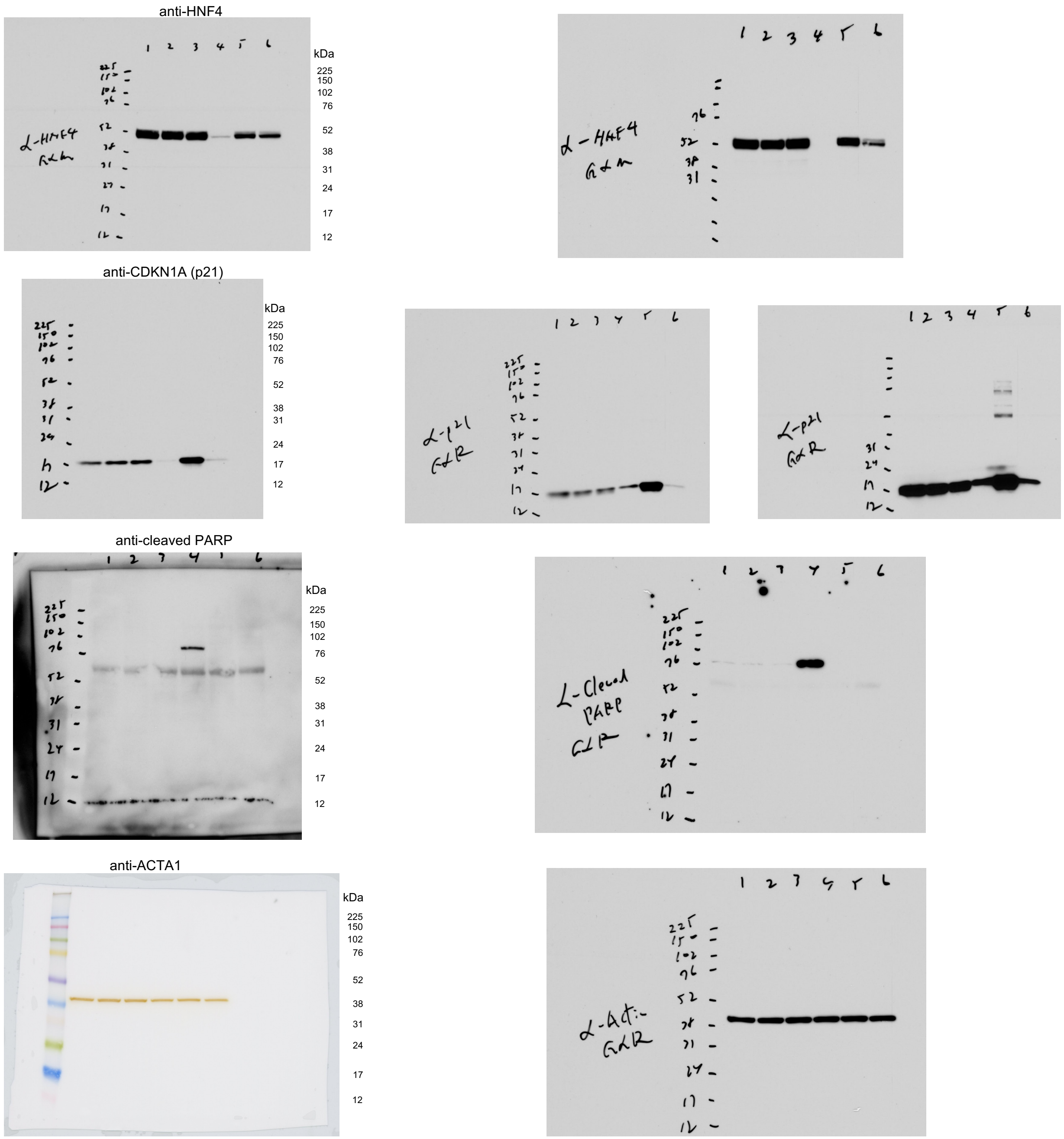


**Supplementary Figure 9. Full size images of western blots using cell extracts from A549 lung carcinoma cells, which are used for Figure 5G.** Two independent experiments were conducted.

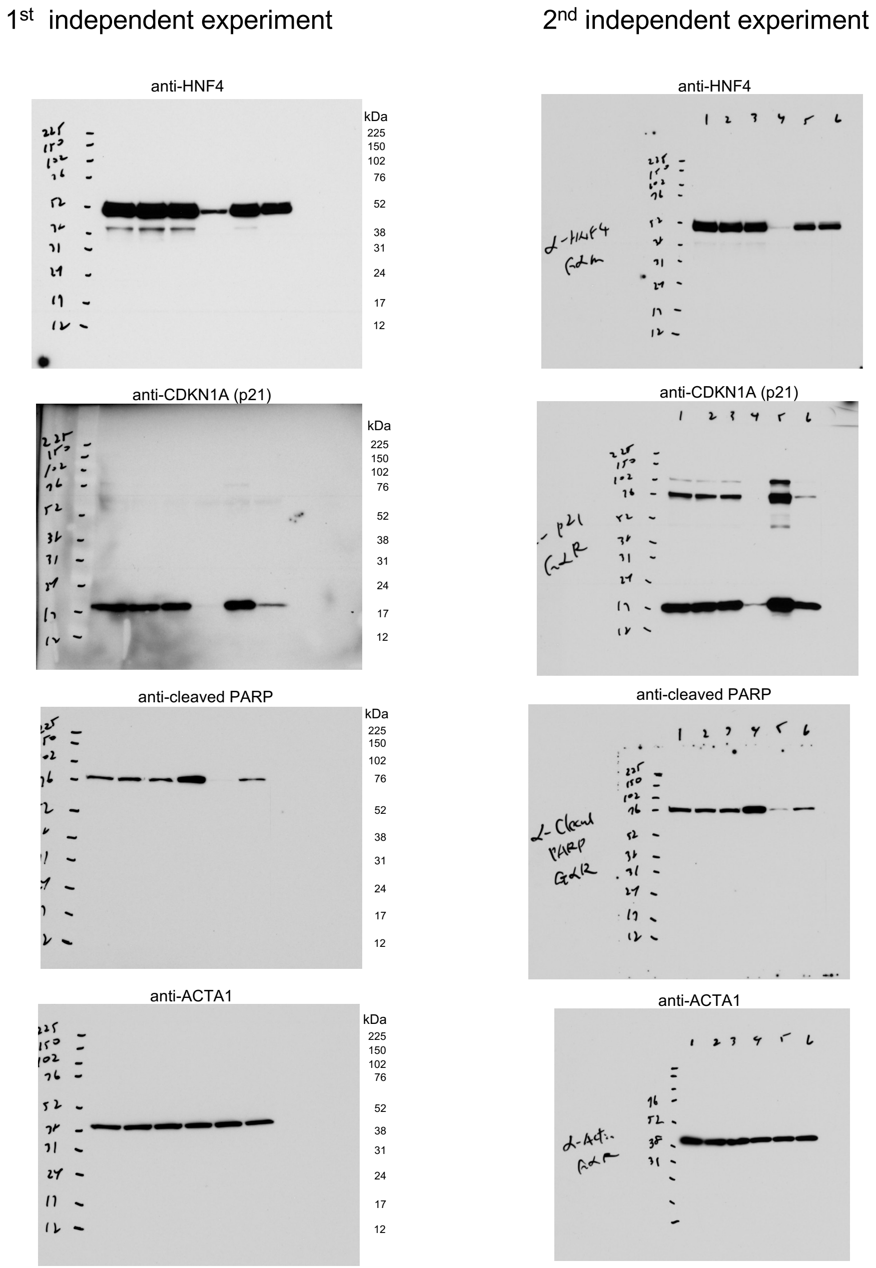


**Supplementary Figure 10. Full size images of western blots using cell extracts from DV-90 lung adenocarcinoma cells, which are used for Figure 5G.** Two independent experiments were conducted.

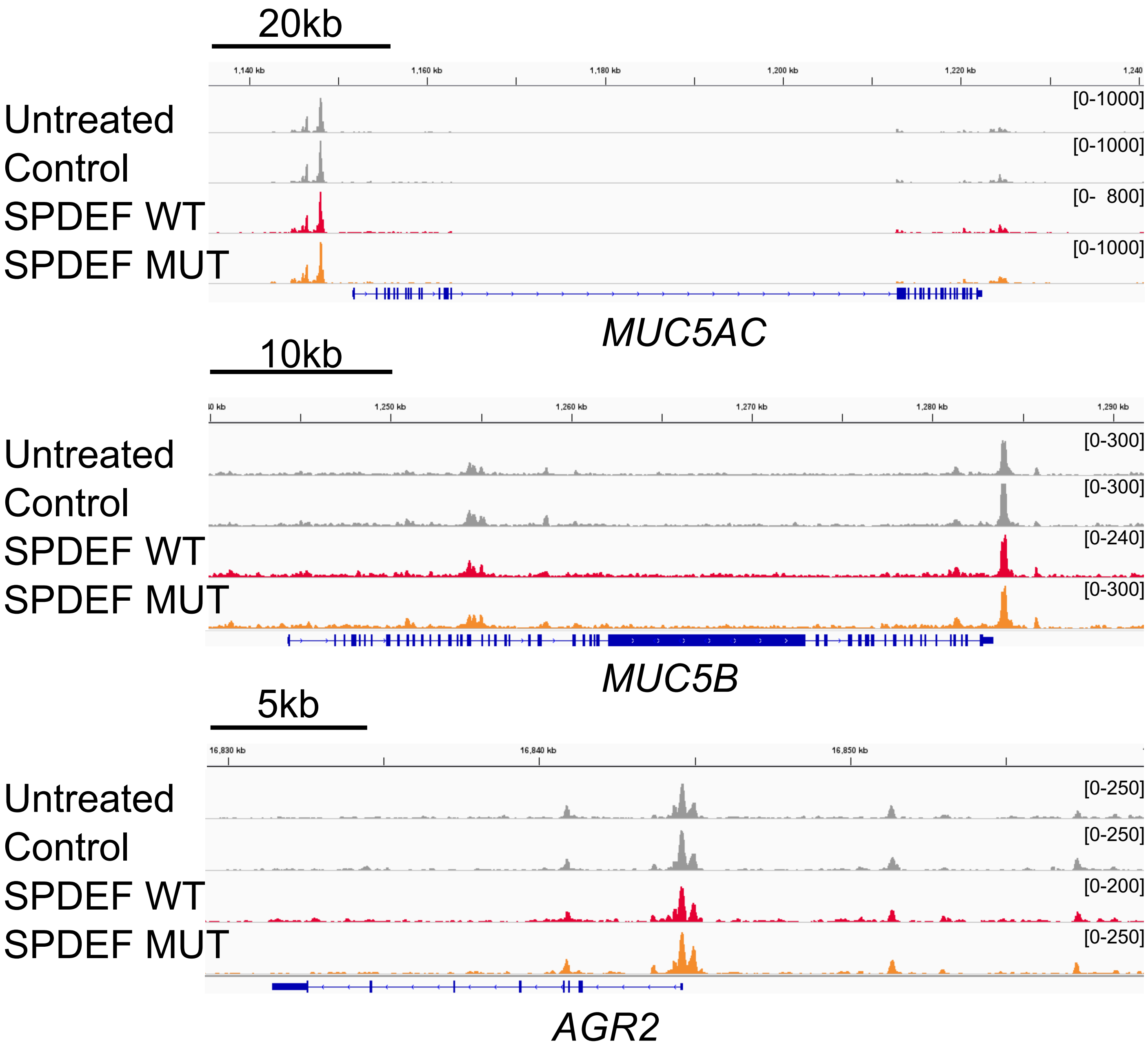


**Supplementary Figure 11. SPDEF does not alter chromatin structure at the mucous gene loci.**
Shown are ATAC-seq peaks indicating that chromatin open regions at *MUC5AC*, *MUC5B* and *AGR2* loci are not affected by the ectopic expression of SPDEF in H292 lung mucoepidermoid carcinoma cell line. Control, SPDEF WT and SPDEF MUT indicate lentiviruses carrying empty, SPDEF wild type and SPDEF G277D, respectively.

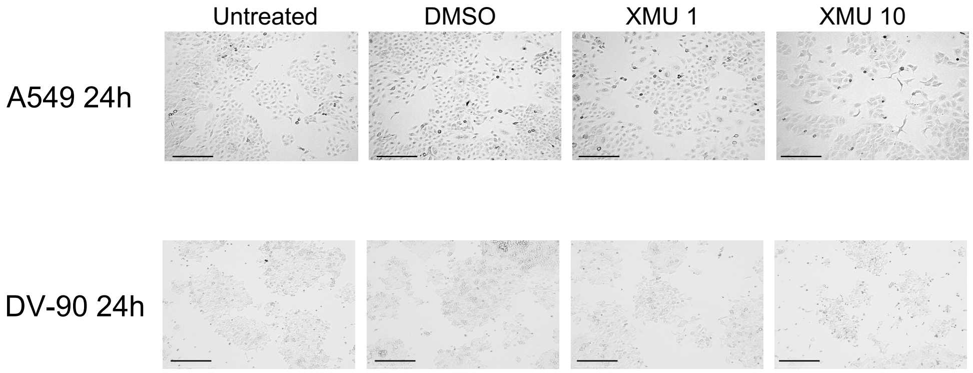


**Supplementary Figure 12. Representative images of A549 lung carcinoma cells and DV-90 lung adenocarcinoma cells treated with XMU-MP-1, an MST1/2 inhibitor (YAP/TAZ activator).**
A549 and DV-90 cells that harbor KRAS mutations were treated with XMU-MP-1, an MST1/2 inhibitor (YAP/TAZ activator), that represses the expression of mucous genes. Shown are representative images of the indicated cells that were treated with DMSO (control) or XMU-MP-1 (1 μM or 10 μM) for 24 hours. Bar indicates 275 μM.

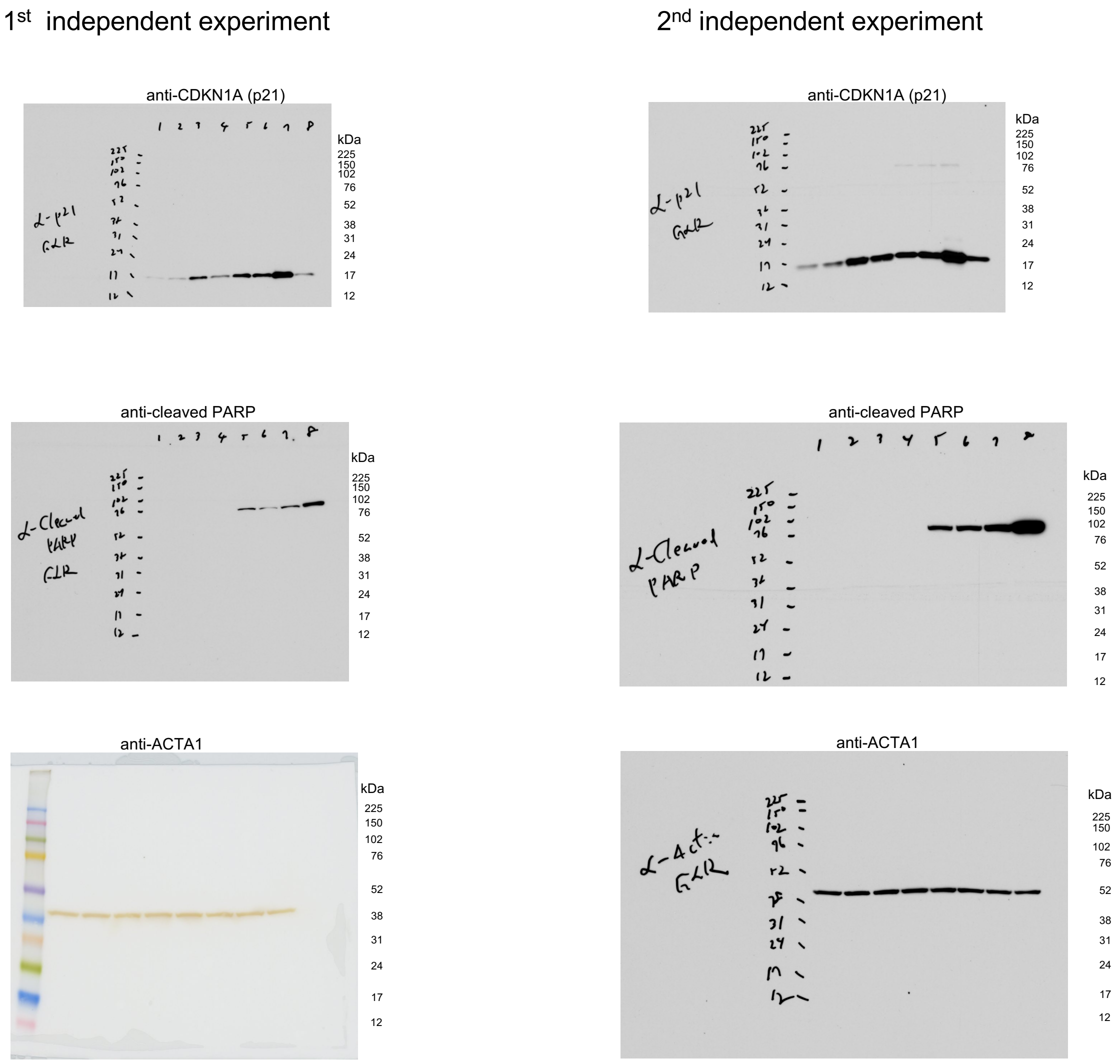


**Supplementary Figure 13. Full size images of western blots using cell extracts from A549 lung carcinoma cells and DV-90 lung adenocarcinoma cells, which are used for Figure 6F.** Two independent experiments were conducted.
